# Perineuronal nets stabilize the grid cell network

**DOI:** 10.1101/796342

**Authors:** Ane Charlotte Christensen, Kristian Kinden Lensjø, Mikkel Elle Lepperød, Svenn-Arne Dragly, Halvard Sutterud, Jan Sigurd Blackstad, Marianne Fyhn, Torkel Hafting

## Abstract

Grid cells are part of a widespread network that supports navigation and spatial memory. Stable grid patterns appear late in development, in concert with extracellular matrix aggregates termed perineuronal nets (PNNs) that condense around inhibitory neurons. To reveal the relationship between stable spatial representations and the presence of PNNs we recorded from populations of neurons in adult rats. We show that removal of PNNs leads to lower inhibitory spiking activity, and reduces grid cells’ ability to create stable representations of a novel environment. Furthermore, in animals with disrupted PNNs, exposure to a novel arena corrupted the spatiotemporal relationships within grid cell modules, and the stored representations of a familiar arena. Finally, we show that PNN removal in entorhinal cortex distorted spatial representations in downstream hippocampal neurons. Together this work suggests that PNNs provide a key stabilizing element for the grid cell network.

## Introduction

Spatially tuned neurons in the hippocampus and entorhinal cortex are key units for navigation and spatial memory. Neurons in the medial entorhinal cortex (MEC) represent information such as self-location (*1*), the direction of the animal’s head (*2*), proximity to geometric borders (*3*), speed (*4*), and possibly the distance traveled by the animal (*5*). The precise position of the animal can be decoded from the activity of grid cell ensembles, where each cell has multiple firing fields forming a characteristic hexagonal pattern spanning the entire surface of the area visited by the animal (*6*).

Grid cells provide input to the hippocampus (*7*) and are assumed to be the primary determinant of hippocampal place cell firing (*3, 5, 8*). However, this notion has been challenged by the fact that place cells appear before grid cells during development (*9, 10*). In rodents, grid cell firing patterns emerge around postnatal day 16-18 (P16-P18) and transition over time from being unstable and non-periodic to highly regular, reaching adult level grid scores around P28-P34 (*9, 11*). Once established, the periodic spiking pattern of grid cells is remarkably stable when animals revisit the same environment. Local perturbations of different cell types in MEC cause changes to grid field spike rates or an increased number of out-of-field spikes, but the location of the fields remains unaltered (*12–14*). Furthermore, when placed in a different environment, neighboring populations of grid cells show a coherent shift of grid fields (*15*) that retains their relative spatiotemporal relationships (*16*). In contrast the hippocampal place cells remap by independently and unpredictably changing their activity. This suggests that the mature grid cell network is more hardwired and less plastic than the hippocampal place cell network.

The network that controls grid cell spiking is not yet fully understood, but recurrent inhibition appears to be fundamental to the specific activity of grid cells. Stellate cells, one of the two principal cell types that display grid cell firing (*7*), are connected via parvalbumin expressing (PV^+^) inhibitory interneurons (*17, 18*). PV^+^ cells account for about 50% of the inhibitory cells in MEC (*19*), making them the main inhibitory mediator in the local grid cell network. In sensory cortex, PV^+^ cells play a central role in shaping the excitatory activity of principal neurons by regulating the onset of periods of high synaptic plasticity (*20*). Whether PV^+^ inhibitory neurons in MEC play a similar role for development of the grid cell network remains elusive.

A hallmark of maturing PV^+^ cells is that aggregates of specialized extracellular matrix, called perineuronal nets (PNNs), condense on the cell soma and proximal dendrites, leaving openings only for synaptic connections (*21*). PNNs are believed to help stabilize the activity of PV^+^ cells by supporting synaptic integrity and limiting synaptogenesis, in addition to supporting PV^+^ cell physiology (*22*). By the time PNNs in sensory cortex are fully mature, plasticity in the local network is strongly reduced. However, juvenile levels of plasticity can be reinstated by experimentally removing PNNs in adult animals, which both increases structural plasticity and reduces inhibitory spiking (*23–25*). Interestingly, the timeline for maturation of PNNs in MEC coincides with the timeline for the development of grid cell firing (*11, 26*). This co-occurrence suggests that grid cell activity could be shaped during the developmental period with high levels of plasticity, and that the later presence of PNNs ensures stability of established synaptic connections, and thus maintains the integrity of the network and the spatiotemporal relationship between grid cells.

To test if PNNs support the stability of the grid cell network, we experimentally disrupted PNNs in MEC of adult rats and recorded from single units while animals explored a familiar arena or a novel environment for the first time. We observed reduced inhibitory spiking activity when PNNs were removed, and grid cell displayed reduced spatial specificity and spatial information in the familiar environment. We replicated these findings in a simulated network with altered synaptic weights that gave rise to reduced inhibition. This is in line with the role for inhibitory neurons in shaping grid cell activity. When the MEC network was challenged with new information by letting animals explore a novel environment, both the spatial correlations of grid cells and their pairwise temporal correlations were significantly reduced in animals lacking PNNs. This indicates that the novel place code remained unstable and the spatiotemporal relationship between grid cells was impaired. The exposure to a novel environment also destabilized the subsequent representation of the familiar arena, suggesting that PNNs are important for maintaining consistent grid cell representations when a previous environment is revisited.

Finally, we recorded place cells from hippocampal area CA1 when PNNs were removed in MEC. Our data shows that the local changes we observed in MEC were also reflected in place cell coding, supporting the idea that the stability and high spatial specificity of grid cell representations are necessary to provide accurate spatial information to place cells.

Together, our data shows that the presence of PNNs ensures precise spatial and temporal coding needed to maintain the network configuration of grid cell and place cell circuits.

## Results

### Degrading PNNs destabilizes synaptic connections and reduces spiking activity of neurons in MEC

The dense expression of PNNs in MEC of adult rats shows an almost complete overlap with expression of PV and mainly enwraps PV^+^ cell soma and proximal dendrites (Fig. 1a). We used local injections of the bacterial enzyme Chondroitinase ABC (chABC) to degrade PNNs, and verified the efficiency in histology sections by labeling of *Wisteria floribunda* agglutinin (WFA)-positive PNNs and 6-sulfated unsaturated disaccharides (3B3 “stubs”) that are left after the degradation process. Staining for WFA and 3B3 stubs clearly delineated the area affected by chABC (Fig. 1b).

**Figure 1:**
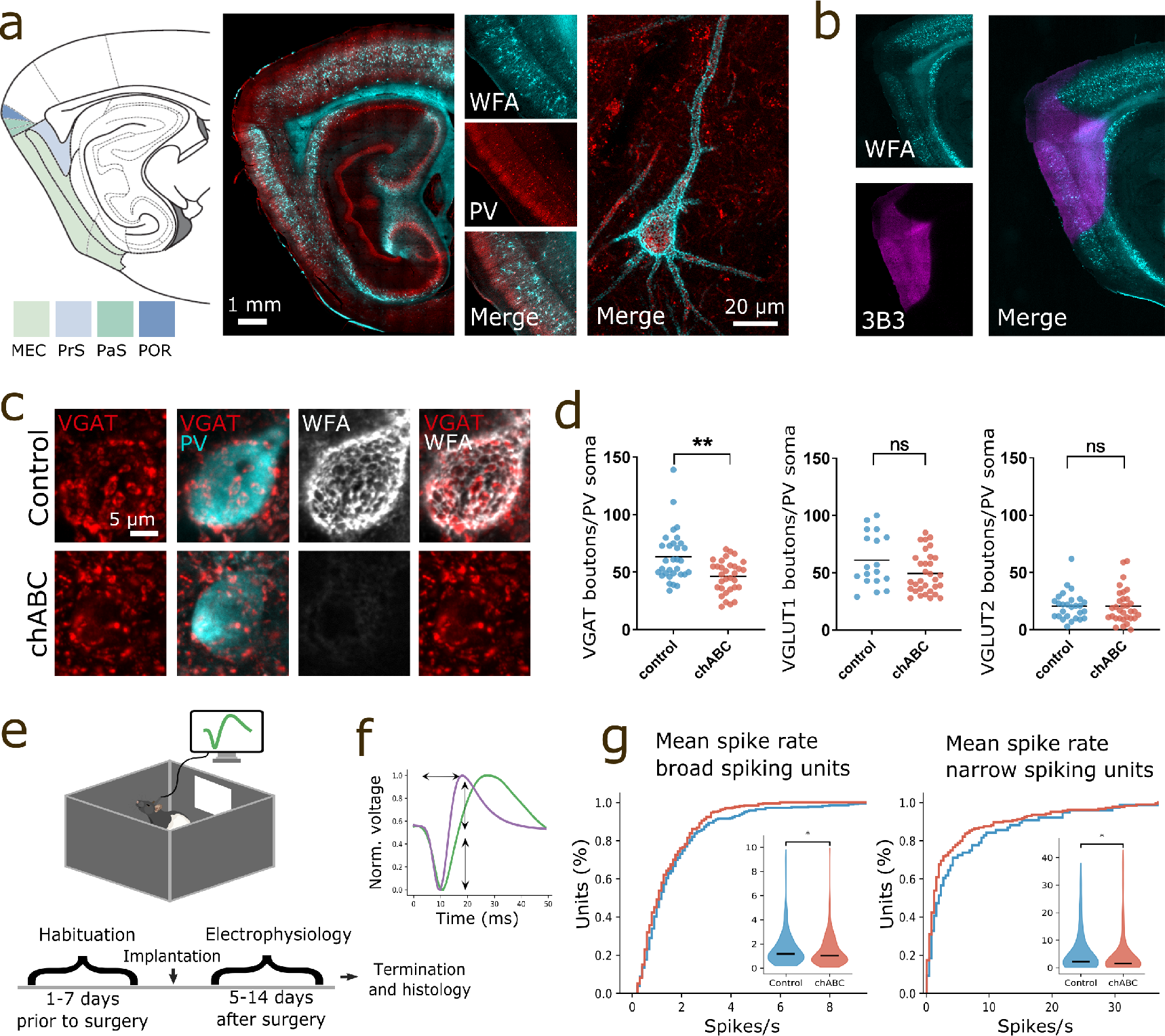
PNNs are highly expressed in MEC where they stabilize synaptic connections and inhibitory neuron spiking activity. **a**) Color coded sagittal drawing outlining parahippocampal areas. Sagittal section from rat brain stained with *Wisteria floribunda* agglutinin (WFA) to label PNNs (cyan) and anti-parvalbumin (PV) antibody to label PV^+^ interneurons (red). PNNs are strongly expressed in MEC and parasubiculum (PaS), while there is weaker staining in presubiculum (PrS) and postrhinal cortex (POR). The overlap between PNNs and PV^+^ neurons is high (large outline). PNNs in MEC enwrap the cell soma and large parts of proximal dendrites (small outline). (Schematic drawing adapted from Paxinos rat brain atlas). **b**) Sagittal section labeled with WFA (cyan) and anti-chondroitin sulfate (3B3) antibody (magenta) to label 6-sulfated unsaturated disaccharide (C-6-S) stubs left from enzymatic degradation of PNNs by chondroitinase ABC (chABC). Intact PNN staining is greatly reduced after chABC injection and C-6-S stubs shows the full extent of the injection area. **c**) VGAT expressing puncta on a PV^+^ cell in MEC five days after injection of aCSF (control, top panel) or chABC (lower panel). **d**) The number of VGAT expressing puncta on PV soma is reduced after local chABC treatment in MEC (VGAT: *p* = 0.002, *n* = 31/32 cells (Control/chABC), mean ± s.e.m.; control 63.29 ± 4.038; chABC 46.34 ± 2.48; VGLUT1: *p* = 0.078, *n* = 18/32, mean ± s.e.m.; Control 60.94 ± 5.48; chABC 49.44±3.13; VGLUT2: *p* = 0.67, *n* = 27/32, mean ± s.e.m.; Control 20.54±2.64; chABC 20.74±2.35. 3 animals, Mann-Whitney U test). **e**) Illustration of open field recording setup and timeline for experiments. **f**) Broad (green) and narrow (purple) spiking units are separated based on the following waveform properties; peak to trough time and half width of amplitude (Fig. S2). **g**) From left: Distribution of mean spike rates for broad -and narrow spiking units from controls and chABC treated rats. Both broad spiking units and narrow spiking units showed reduced spike rates in animals with disrupted PNNs (mean ± s.e.m.; Control broad spiking 1.68 ± 0.09; chABC broad spiking 1.40 ± 0.07, *p* = 0.025, Control *n* = 293, chABC *n* = 278; Control narrow spiking 5.6 ± 0.88. chABC narrow spiking 4.86± 0.62, *p* = 0.031, Control *n* = 78, chABC *n* = 187, Mann-Whitney U test). Violin plot shows min to max and median (large black line). *ns* = *notsignificant*, \**p* < 0.05, \*\**p* < 0.01, \*\*\**p* < 0.001, **** *p* < 0.0001.

PNNs are hypothesized to limit plasticity by stabilizing synaptic connections and facilitating the high spiking activity of PV^+^ cells (*27–29*). To test how PNNs affect synaptic stability in MEC, we first degraded PNNs unilaterally using chABC. Five days after chABC treatment, we quantified synaptic boutons contacting the soma of PV^+^ cells via immunostaining. Because environmental enrichment such as exploration of novel arenas or changes to the housing environment can affect neural plasticity (*30*), animals were kept in their home cage throughout the experiment to isolate the effect of PNN degradation.

In the control hemisphere, we found that a large fraction of inputs to PV^+^ cell somas were positive for the inhibitory presynaptic marker VGAT (Fig. 1c), while only a few were positive for VGLUT2 (Fig. S1). Removing the PNNs caused a reduction in the number of VGAT expressing puncta onto PV^+^ cells (*p* = 0.002, Mann-Whitney U test, 3 rats, *n* aCSF =31 cells, *n* chABC =32 cells). No significant changes were observed for excitatory VGLUT 1 and 2 puncta (Fig. 1d), although there was a clear tendency for reduction in VGLUT 1 puncta onto PV^+^ cells. To investigate the functional role of PNNs, we next recorded single unit activity from MEC using bilaterally implanted tetrodes in 13 rats (7 of which were treated with chABC) while the animals explored an open field arena (Fig. 1e). All units recorded in a familiar environment were separated into narrow- and broad spiking based on waveform properties (Fig. 1f, Fig. S2) (*31, 32*). Narrow spiking units are putative inhibitory neurons of which PV^+^ cells constitute the largest group (*19, 33*), while broad spiking units are putative excitatory neurons. A total of 283 units were identified as broad spiking in the control group and 278 in the chABC group. The number of narrow spiking units were 76 (20%) in the control group and 187 (40%) in the chABC group. Both broad- and narrow spiking units showed decreased mean spike rates in chABC treated animals (Fig. 1g) These results are in line with recordings from the visual cortex (*24*).

### Spatial specificity and information is reduced in familiar environments

Grid cells are widely thought to correspond to layer II principal cells in MEC (*18, 34*). Layer II principle cells, and in particular stellate cells display profound inter-connectivity with PV^+^ neurons, contributing to the sharp receptive fields and low out-of-field noise of grid cells (*12, 17, 18, 35*). To examine if PNNs are required for grid cell spiking activity, we classified grid cells using a dual criteria based on gridness score and spatial information analysis. Units that had scores above the 95th percentile for both gridness scores and spatial information, generated from shuffling each units’ own spikes, were classified as grid cells. If a unit was recorded more than once, only the first recording was included in statistical analysis of grid cell spiking properties (Table 1). From this, we identified a total of 840 unique units, of which 23% displayed grid cell spiking activity in the control group (86 out of 373 units), and 14% in chABC treated animals (63 out of 467 units).

**Table 1:** Firing properties of grid cells in a familiar environment for control and chABC groups (mean ± s.e.m. (n), Mann Whitney U test (U, p) and Permutation resampling (diff, p)).

|  | Control | chABC | MWU | PRS |
| --- | --- | --- | --- | --- |
| Average rate | 1.98 ± 0.18 (86) | 1.91 ± 0.17 (63) | 2695.00, 0.959 | 0.13, 0.605 |
| Gridness | 0.62 ± 0.03 (86) | 0.60 ± 0.03 (63) | 2812.00, 0.694 | 0.07, 0.371 |
| Max rate | 11.29 ± 0.76 (86) | 8.33 ± 0.72 (63) | 3385.00, 0.009 | 2.39, 0.012 |
| Information rate | 0.53 ± 0.05 (86) | 0.29 ± 0.03 (63) | 3688.00, 0.000 | 0.17, 0.005 |
| CV of interspike interval | 2.02 ± 0.06 (86) | 1.62 ± 0.05 (63) | 3975.00, 0.000 | 0.31, 0.000 |
| In-field mean rate | 3.80 ± 0.29 (86) | 3.12 ± 0.24 (63) | 3059.00, 0.179 | 0.10, 0.501 |
| Out-field mean rate | 1.10 ± 0.12 (86) | 1.28 ± 0.15 (63) | 2395.00, 0.228 | 0.10, 0.198 |
| Burst event ratio | 0.09 ± 0.01 (86) | 0.04 ± 0.01 (63) | 4148.50, 0.000 | 0.06, 0.000 |
| Specificity | 0.57 ± 0.02 (86) | 0.45 ± 0.02 (63) | 3796.00, 0.000 | 0.13, 0.000 |

In animals treated with chABC we observed more dispersed spiking within and at the edges of grid fields (Fig. 2a and b), but this did not lead to reduction in gridness scores (Fig. 2c). Pharmacogenetically reducing activity in PV^+^ neurons lead to increased out-of-field activity in grid cells (*12*). To test if similar effects could be seen after removal of PNNs, we identified firing fields for each grid cell (Fig. S3) and calculated mean firing rate inside and outside of fields. We found a weak but consistent effect of PNN removal, where grid cells from chABC treated rats showed reduced firing rates inside grid fields and increased firing rates outside grid fields (Table 1). The combination of these effects lead to a prominent decrease in specificity of grid cells in animals with disrupted PNNs (Fig. 2d). To account for effects of an arbitrary definition of grid fields, we also calculated the spatial information content of each grid cell, a measure that is independent of any predefined firing fields (*36*). The spatial information was significantly reduced in grid cells from animals treated with chABC (Fig. 2e). In addition, we found changes in the temporal spiking properties of grid cells. Maximum firing rate was significantly reduced (Fig. 2f), along with the fraction of bursting events (Fig. 2g). This was accompanied by a reduction in spiking variability, measured by the coefficient of variation (CV) of interspike intervals (ISI) (Fig. 2h, Fig. S4).

**Figure 2:**
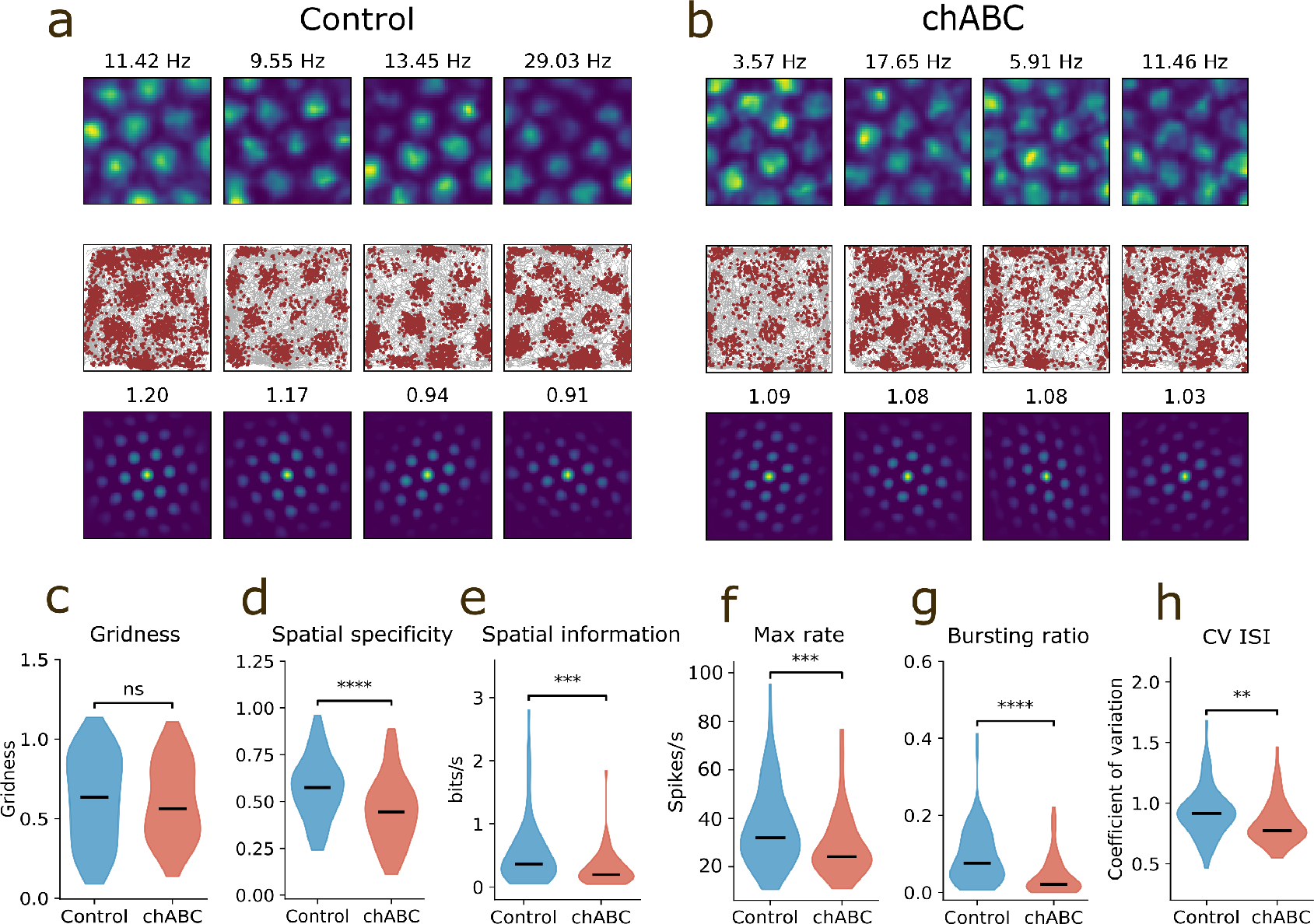
Grid cells show reduced spatial coding after PNN removal. **a-b**) Example grid cells from control and chABC treated rats. Color coded rate maps (top), trajectory maps with spikes superimposed (middle) and spatial autocorrelation maps (bottom) for four grid cells in controls (a) and four grid cells in chABC treated rats (b). The number above the rate map is maximum firing rate and the number above the autocorrelation map is gridness score. **c**) Removing PNNs did not reduce gridness in the familiar environment (mean ± s.e.m.; Control 0.62 ± 0.03; chABC 0.60 ± 0.03, *p* = 0.694). **d**) The dispersed spiking outside grid cell fields caused the spatial specificity of grid cells to decrease (mean ± s.e.m.; Control 0.57 ± 0.02; chABC 0.40 ± 0.02, *p* < 0.0001), in addition to reducing the **e**) spatial information (mean ± s.e.m.; control 0.53 ± 0.05; chABC 0.29 ± 0.03, *p* = 0.000). **f**) Maximum firing rate was reduced (mean ± s.e.m.; Control 36.40± 1.82; chABC 28.62 ± 1.68, *p* = 0.000). **g**) Grid cell bursting ratio was measured as the number of bursting events divided by the number of single-spike events. Rats with disrupted PNNs showed a large reduction in bursting event ratio (mean ± s.e.m.; Control 0.09±0.01; chABC 0.04 ± 0.01, *p* < 0.001). **h**) Spiking variability was measured as the coefficient of variation (CV) of interspike intervals (ISI). The method used here calculates CV of ISI using only spikes from passes though grid fields (Fig. S4), (mean ± s.e.m.; Control 0.93 ± 0.02; chABC 0.83 ± 0.02, *p* = 0.001). Violin plots show min to max and median (black line). Width of graph corresponds to number of samples for each value. *n* Control =86, *n* chABC =63, all tests are Mann-Whitney U tests. *p < 0.05, **p < 0.01, ***p < 0.001, **** *p* < 0.0001.

### Grid cell representations are impaired in novel environments

Previous work from other cortical areas has demonstrated that PNN removal dramatically increases neural plasticity. However, the removal does not cause major disturbances to normal network function unless the system is required to encode new information. In the visual cortex, properties such as receptive fields and visual acuity are intact when the PNN is removed, but undergo dramatic changes if vision is occluded from one eye (*23, 24*). Since we observed only subtle effects on grid cell spiking properties in the familiar environment when PNNs were removed, we wanted to test how the grid cell network responded to encoding new information. We introduced the animals to a novel environment which cause grid cells to remap (shift their grid map) (*15*), expand and become spatially unstable (*37*), an effect that is reversed when animals explore the environment to the point where it becomes familiar.

When stable grid cells could be recorded in the familiar environment, the animals were introduced to a similar box in a novel room and allowed to explore for 3 × 20 minutes (Fig. 3a).

**Figure 3:**
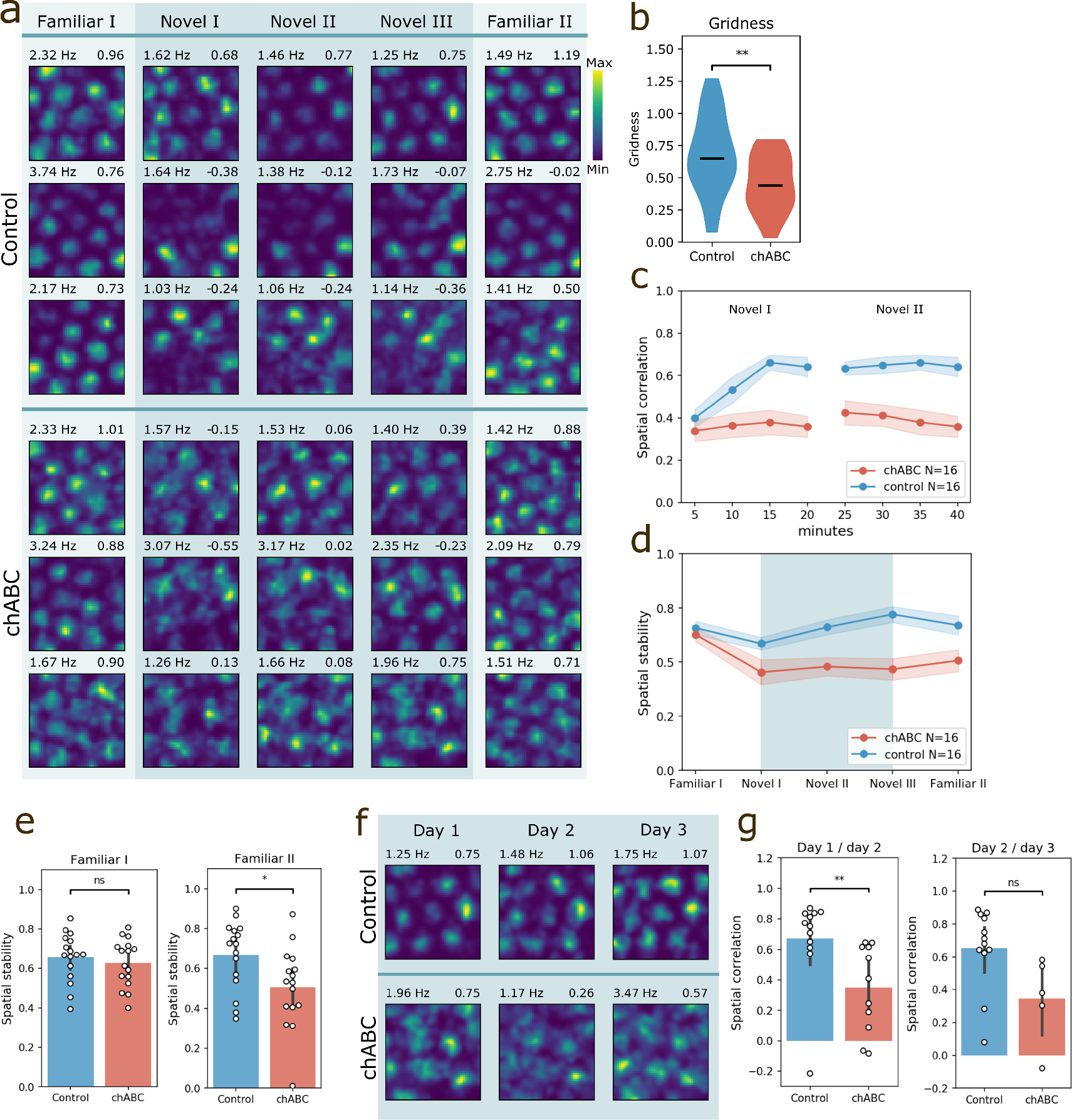
Grid cells from chABC treated animals show reduced spatial correlations in a novel environment. **a**) Rate maps of five consecutive 20 minutes recording sessions from six individual grid cells, recorded in control animals (top) and chABC animals (bottom) during a novel environment experiment. Color code indicates spike rate. Rows correspond to individual units and columns correspond to environment. Maximum spike rate (left) and gridness (right) is denoted above each rate map. **b**) Gridness score is reduced in the novel environment in animals treated with chABC ((mean ± s.e.m.; Control 0.68 ± 0.30; chABC 0.46 ± 0.22; *p* = 0.004, Mann-Whitney U test). **c**) Spatial correlation measured for blocks of five minute recordings in the novel environment sessions I & II, measured against the novel environment session III. Only units reaching gridness threshold in Familiar I and that could be identified in all five recording sessions were included. Animals treated with chABC showed reduced spatial correlations in the novel environment (mean ± s.e.m.; Control 0.59 ± 0.03; chABC 0.37 ± 0.04, main effect of group: *F*(*1, 30*) = 14.22, *p* = 0.0007, interaction: *F*(*7, 210*) = 4.905, *p* < 0.0001, Control *n* = 16; chABC *n* = 16, two-way repeated measures ANOVA with group and time as factors). **d**) Spatial stability of the same units as in **c**, measured by the within-trial spatial correlation of grid cell rate maps (0-10 min correlated with 10-20 min). Spatial stability decreased from Familiar I to Novel I in the chABC treated group, but not in the control group (mean ± s.e.m.; Control Familiar I 0.66 ± 0.03; Control Novel I 0.59 ± 0.03, *p* = 0.086; chABC Familiar I 0.63 ± 0.03; chABC Novel I 0.45 ± 0.06, *p* = 0.015, Mann-Whitney U test). **e**) Spatial stability was similar for both groups in Familiar I (mean ± s.e.m.; Control 0.66 ± 0.03; chABC 0.63 ± 0.03, *p* = 0.629). However, after exploring the novel environment, chABC animals showed reduced spatial stability upon returning to the familiar environment (Familiar II), (mean ± s.d.; control 0.67 ± 0.04; chABC 0.51 ± 0.05, *p* = 0.018, Control *n* = 16; chABC *n* = 16, Mann Whitney U test). **f**) Rate maps of two grid cells when introduced to the novel environment for three consecutive days. **g**) Spatial correlation of grid cells from chABC treated animals was decreased in the novel environment when comparing day 1 with day 2 (mean ± s.e.m.; control 0.67 ± 0.09, *n* = 13; chABC 0.35 ± 0.09, *n* =11, *p* = 0.004, Mann-Whitney U test). Rate map from the third novel session (Novel III) on the first day was used for correlations with day 2. *ns* =not significant, *p < 0.05, **p < 0.01, ***p < 0.001, **** *p* < 0.0001

While the grid cells’ gridness scores was similar between groups in the familiar environment, introduction to a novel environment lead to a reduction in gridness scores in animals with disrupted PNNs. When averaging all three sessions in the novel environment, control animals showed gridness scores similar to the familiar environment, while the chABC treated group had substantially reduced gridness scores in the novel versus familiar environment (Fig. 3b, Table 2). We then tested how fast newly formed grid maps adapted a stable spatial spiking pattern by correlating rate maps for every five minutes of the first two novel environment recording sessions (Novel I and II) with the rate map from the last 20 minutes in the novel room (Novel III) (a measure we abbreviate to *spatial correlation*). Only units that were categorized as grid cells in familiar I and could be followed throughout all recording sessions were included in this analysis. Animals injected with chABC in MEC showed significantly lower spatial correlations in the novel environment (main effect of group: *F*(*7, 210*) = 4.905, *p* < 0.0001; interaction: *F*(*7, 21*) = 4.91, *p* < 0.0001, Control *n* = 16, chABC *n* = 16, Two-way repeated measures ANOVA with group and time as factors). Correlations were lower in all but the first two periods, 0 to 5 minutes and from 5 to 10 minutes (Šidák’s multiple comparisons post hoc test, Table S1). Animals treated with chABC did not reach spatial correlation levels comparable to control animals within 60 min of exploring the novel environment (Fig. 3c).

**Table 2:** Spatial stability of grid cell representations. Upper panel compares groups while lower shows in-group comparisons. Values are mean ± s.e.m. (n), Mann-Whitney (MWU) (U, p) and Permutation Resampling (PRS) (diff, p).

| Between groups | Control | chABC | MWU | PRS |
| --- | --- | --- | --- | --- |
| Gridness | $0.68 \pm 0.30$ (16) | $0.46 \pm 0.22$ (16) | 606.00, 0.004 | 0.23, 0.002 |
| Spatial stability Familiar I | $0.66 \pm 0.03$ (16) | $0.63 \pm 0.03$ (16) | 139, 0.692 | 0.02, 0.629 |
| Spatial stability Familiar II | $0.67 \pm 0.04$ (16) | $0.51 \pm 0.05$ (16) | 191, 0.018 | 0.19, 0.016 |
| Familiar I vs Familiar II | $0.63 \pm 0.08$ (16) | $0.57 \pm 0.08$ (16) | 152, 0.376 | 0.08, 0.162 |
| Novel day1 vs Novel day2 | $0.67 \pm 0.09$ (12) | $0.35 \pm 0.09$ (11) | 113, 0.004 | 0.35, 0.006 |
| Novel day2 vs day3 | $0.66 \pm 0.07$ (12) | $0.35 \pm 0.12$ (5) | 52, 0.023 | 0.32, 0.052 |
| Within groups | Control | chABC |  |  |
| Spatial stability Familiar I | $0.66 \pm 0.03$ (16) | $0.63 \pm 0.03$ (16) | | |
| Spatial stability Novel I | $0.59 \pm 0.03$ (16) | $0.45 \pm 0.06$ (16) | | |
| PRS | 0.13, 0.070 | 0.21, 0.017 |  |  |
| MWU | 174, 0.086 | 193, 0.015 |  |  |

We next aimed to test the *spatial stability* of the grid maps during each session by calculating the within-trail spatial correlation. To calculate spatial stability, the rate map from the first 10 minutes of a recording session was correlated against the rate map from the last 10 minutes of the session. Within-trail stability was significantly reduced from the familiar environment to the novel environment in the chABC group but not in the control group, and remained reduced throughout all sessions in the novel environment (Fig. 3d).

Interestingly, while the spatial stability in the first familiar session did not differ between the two groups, the brief exposure to the novel arena affected the subsequent coding of the familiar environment in chABC treated animals, as shown by reduced spatial stability within the familiar II trial (Fig. 3e).

In our experimental design, rats were re-introduced to the same “novel” environment for three consecutive days (Fig. 3f). Animals treated with chABC continued to show reduced spatial correlation when comparing the last novel environment recording session on day 1 (Novel III) with the novel environment recording session on day 2. This indicates that novel spatial maps were not properly stabilized during the 60 min exploration time on day 1. There was also a strong tendency towards reduced spatial correlation in the novel environment from day 2 to day 3, but the units identified as the same from day 1 to day 3 in the chABC treated group were too few to ensure proper statistical power (Fig. 3g).

### PNN removal alters theta oscillations

Local field potentials (LFPs) in MEC show strong oscillations in the theta frequency range (6-12 hz) when rats are moving. These oscillations modulate grid cell spiking and organize neuronal activity into temporal windows that is believed to be important for episodic memory formation and plasticity (*38, 39*). PV^+^ neurons are vital for producing synchronized activity in local networks and for organizing information transfer between brain areas (*40, 41*). Hence, changes in PV^+^ cell function and inhibitory plasticity could have a widespread effect on the synchronized activity of the MEC network. To address this we analyzed the LFPs recorded from MEC of 14 animals (of which 7 were injected with chABC). We found that LFPs were altered in the theta range of chABC treated rats (Fig. 4a), but we observed no change in other frequencies (data not shown). The average power of theta oscillations was stronger and the peak frequency decreased in the chABC treated rats in both novel and familiar environments (Table 3).

**Table 3:** Theta frequency and power in familiar and novel environments. Correlation between running speed and power or frequency: power score and frequency score. Showing mean ± s.e.m. (n), Mann-Whitney (MWU) (U, p) and Permutation Resampling (PRS) (diff, p).

|  | Control | chABC | MWU | PRS |
| --- | --- | --- | --- | --- |
| Power familiar | -6.41 ± 0.21 (148) | -4.56 ± 0.22 (75) | 2832, 0.000 | 1.94, 0.000 |
| Power novel | -7.15 ± 0.38 (33) | -4.74 ± 0.21 (24) | 126, 0.000 | 2.37, 0.000 |
| Frequency familiar | 8.44 ± 0.04 (148) | 8.31 ± 0.04 (75) | 6356, 0.075 | 0.12, 0.003 |
| Frequency novel | 8.56 ± 0.11 (33) | 8.14 ± 0.07 (24) | 558, 0.009 | 0.30, 0.033 |
| Power score familiar | 0.17 ± 0.01 (148) | 0.32 ± 0.02 (75) | 2800, 0.000 | 0.17, 0.000 |
| Power score novel | 0.13 ± 0.01 (33) | 0.29 ± 0.02 (24) | 102, 0.000 | 0.17, 0.000 |
| Frequency score familiar | 0.14 ± 0.01 (148) | 0.20 ± 0.01 (75) | 3100, 0.000 | 0.06, 0.000 |
| Frequency score novel | 0.12 ± 0.01 (33) | 0.18 ± 0.02 (24) | 200, 0.002 | 0.07, 0.000 |

**Figure 4:**
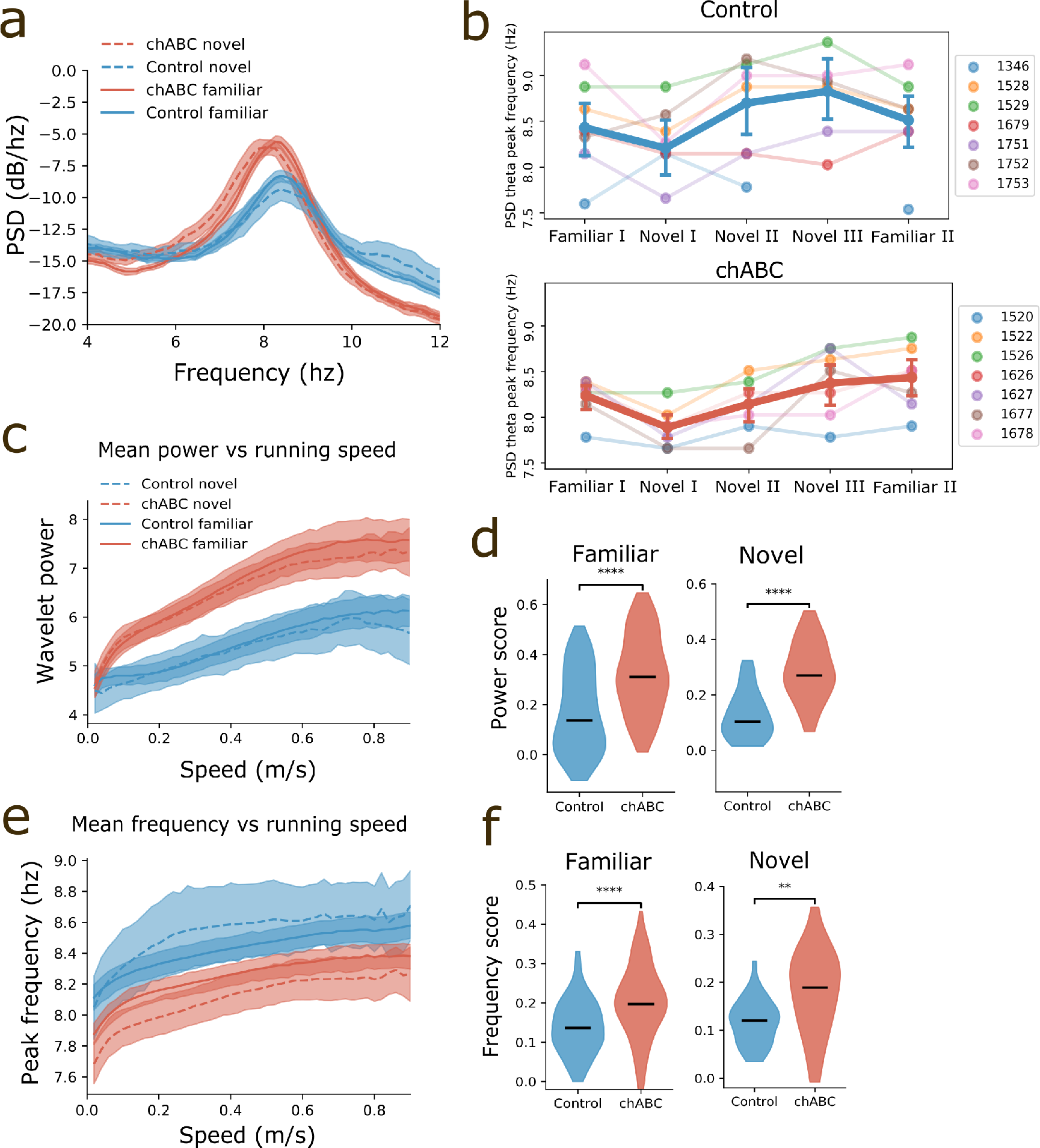
Increased theta oscillations after PNN removal. **a**) Power spectrum of local field potential in MEC shows that theta oscillations in chABC rats have increased power (mean ± s.e.m.; Control familiar 0.27 ± 0.01; chABC familiar 0.37 ± 0.01; Control novel 0.22 ± 0.02; chABC novel 0.34 ± 0.02) and lower peak frequency (mean ± s.e.m.; Control familiar 8.44 ± 0.04; chABC familiar 8.31 ± 0.04; Control novel 8.56 ± 0.11; chABC novel 8.14 ± 0.07) (solid line: familiar, dotted line: novel, blue: control, red: chABC). Shaded area represent 95% confidence interval. **b**) Average theta frequency for all animals during recordings in familiar and novel environments (dark colored line). Note the decrease in theta frequency during first exposure to a novel environment in both groups. Light colored lines correspond to individual animals (left: animal numbers). **c**) The mean theta power was similar when the animal was immobile, but the chABC group showed increased theta power during movement. **d**) Power score shows increased correlation between theta power and running speed in animals treated with chABC for both familiar and novel environments (mean ± s.e.m.; Control 0.17±0.01(*n* = 148); chABC 0.32±0.02(*n* = 75), Control 0.13±0.01(*n* = 33); chABC 0.29±0.02(*n* = 24), respectively). **e**) Theta frequency increased slightly with running speed in both groups, but the peak frequency was lower in the chABC treated group. **f**) Frequency score shows increased correlation between theta frequency and running speed in chABC treated animals for both familiar and novel environment (mean ± s.e.m.; Control 0.14±0.01(*n* = 148); chABC 0.20±0.01(*n* = 75), Control 0.12±0.01(*n* = 33); chABC 0.18±0.02(*n* = 24), respectively).

**Table 4:** Parameters used in the can model.

|  | Excitatory | Control<br>Inhibitory | Increased inhibition<br>Inhibitory | Unit |
| --- | --- | --- | --- | --- |
| $C_m$ | 70.0 | 30.0 | 30.0 | pF |
| $\Delta_T$ | 2.5 | 2.5 | 2.5 | ms |
| $E_L$ | -60.0 | -60.0 | -60.0 | mV |
| $E_{rev,AMPA}$ | | 0.0 | 0.0 | mV |
| $E_{rev,NMDA}$ | | 0.0 | 0.0 | mV |
| $E_{rev,GABA_A}$ | -75.0 | | | mV |
| $\bar{g}_{AMPA}$ | | 0.02 | 0.005 | nS |
| $\bar{g}_{NMDA}$ | | 0.007 | 0.0065 | nS |
| $\bar{g}_{GABA_A}$ | 0.01 | | | nS |
| $\tau_{rise,AMPA}$ | | 1.0 | 1.0 | ms |
| $\tau_{rise,NMDA}$ | | 10.0 | 10.0 | ms |
| $\tau_{rise,GABA_A}$ | 1.0 | | | ms |
| $\tau_{decay,AMPA}$ | | 5.0 | 5.0 | ms |
| $\tau_{decay,NMDA}$ | | 100.0 | 100.0 | ms |
| $\tau_{decay,GABA_A}$ | 5.0 | | | ms |
| $N$ | 10000 | 10000 | 10000 | |
| $V_m$ | -60.0 | -60.0 | -60.0 | mV |
| $V_{peak}$ | -40.0 | -40.0 | -40.0 | mV |
| $V_{reset}$ | -60.0 | -60.0 | -60.0 | mV |
| $V_{th}$ | -50.0 | -45.0 | -45.0 | mV |
| $gL$ | 4.0 | 4.5 | 4.5 | nS |
| $r_{inner}$ | $0.2\pi$ | 0 | 0 | |
| $r_{outer}$ | $0.4\pi$ | $0.15\pi$ | $0.15\pi$ | |
| $p\bar{g}$ | 0.1 | 0 | 0 | nS |
| $p_{rate}$ | 850.0 | 0.0 | 0.0 | Hz |
| $t_{ref}$ | 1.0 | 1.0 | 1.0 | ms |

For both groups the first introduction to the novel environment (Novel I) reduced the theta peak frequency in the power spectral density (PSD) (Fig. 4b), while continued exploration of the novel environment (Novel II and III) lead to a large increase of theta frequency in the control group and a smaller increase in the chABC group.

Theta power and frequency is known to increase with the animals’ running speed (*42, 43*). Hence, to test if the difference in theta between the groups was caused by differences in the animals’ running speed, we measured running speed and calculated a continuous wavelet trans-form, conditioned average theta power and peak frequency in 1 sec time bins of every session. As expected, the peak theta power and frequency had a positive correlation with running speed for both groups (Fig. 4c, e). We found increased theta power in chABC animals for all running speeds in both familiar and novel environments. The only exception was during immobility where the theta power between groups was similar (Fig. 4c). Correspondingly, the frequency of theta was reduced in chABC treated animals relative to controls, regardless of running speed (Fig. 4e). To assess the differences in total correlations between power and running speed we calculated a power score. (Fig. 4d). This was significantly stronger for chABC treated animals in both familiar and novel environments, despite the fact that animals in the chABC group had slightly lower running speeds in novel environments (Fig. S5a). We did the same for theta frequency and running speed (frequency score), where chABC treated animals again showed a significantly higher frequency score (Fig. 4f). Hence, both theta power and theta frequency was more correlated with running speed in animals lacking PNNs, meaning that neither the altered theta frequency or power is well explained by running speed.

### PNN removal causes reduced pairwise stability of grid cells during novel arena exploration

Grid cells that share similar spacing and orientation maintain their relative spatial relationship when the grid map changes in a novel environment (*15, 16, 44*), and the extent of temporal pairwise correlation reflects the degree of spatial overlap. This is indicative of a highly structured and fixed network. Our results suggest that PNN removal increases the potential for plasticity and destabilizes the network in a novel environment. We therefore tested whether this destabilization affected relative spatial and temporal properties within the grid cell population.

We calculated the pairwise spatial and temporal cross-correlation function of grid cells recorded within each experimental paradigm (Familiar I II and Novel I, II, III). We next calculated the correlation of the cross-correlation functions across respective paradigms (for example Familiar I vs Novel I) - if the pairwise spatiotemporal relationship is constant, this correlation should be high. Since we already observed changes in LFP theta, we controlled for theta correlations by removing the band (4-10 Hz) from the firing rate of each neuron.

As expected, control animals showed strong correlations of temporal cross correlations between the familiar and the novel environment (Fig. 5a, c). However, animals treated with chABC showed reductions in pairwise correlations when introduced to a novel environment (Fig. 5b, c), and were significantly less correlated than controls (Fig. 5d). This was also seen when correlating spatial fields (Fig. 5e, f, Fig. S6). Interestingly, chABC animals also showed reduced temporal correlations when returning to the familiar environment (Familiar II) (Fig. 5d). To correct for increased noise deriving from the reduced spatial specificity in the chABC group, we removed all spikes from outside fields and performed the same analysis using only the spikes belonging to grid fields (data not shown). This did not change the results for neither temporal or spatial pairwise correlations.

**Figure 5:**
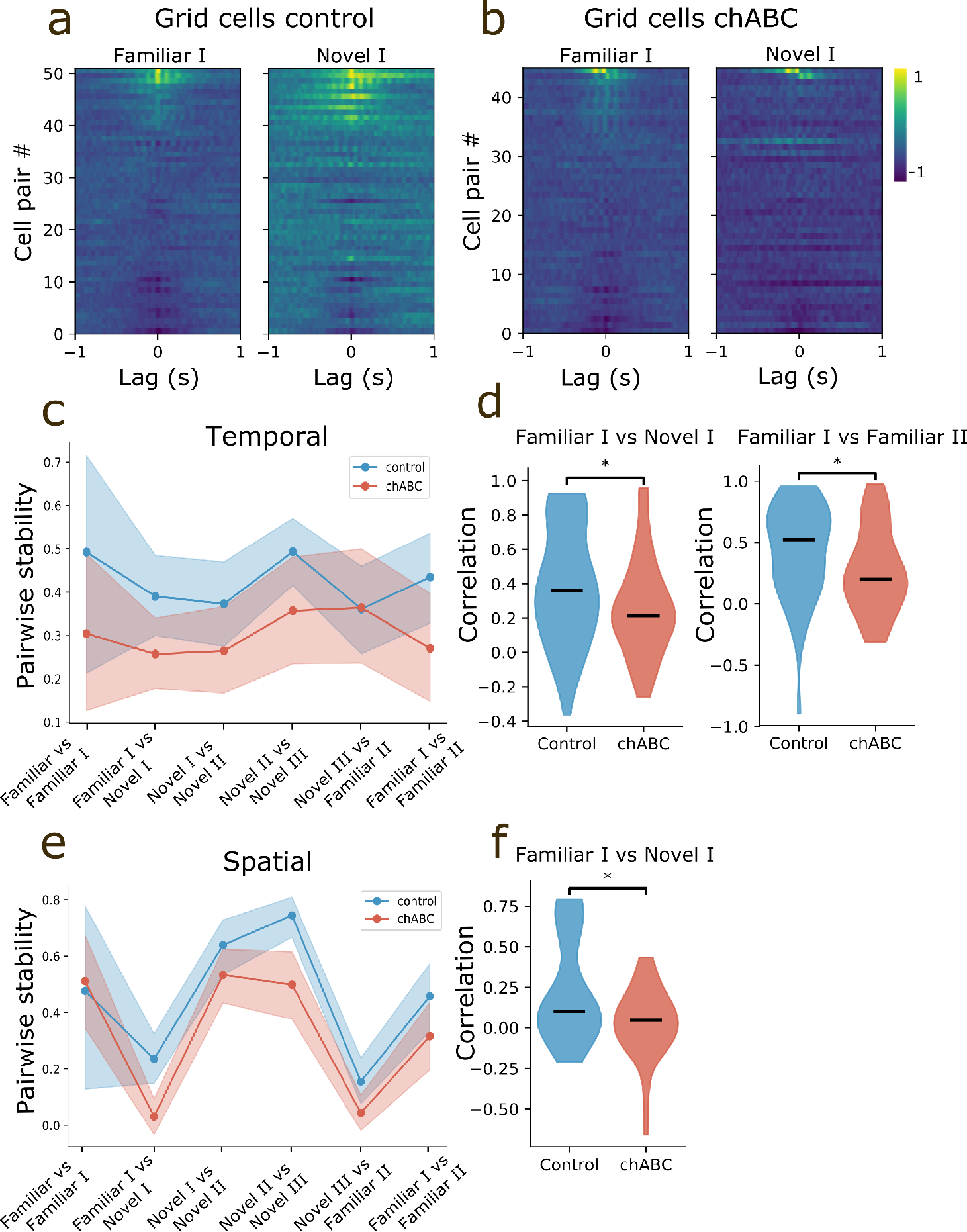
Grid cells are pairwise stable between familiar and novel environments in control, but not in chABC treated animals. **a**) Pairwise temporal cross-correlation of grid cells with brighter color showing stronger correlation. Values range from −1 to 1. Each line represents a cell pair, sorted by maximum value of the central peak. **b**) Same as in **a**, but for chABC treated animals. **c**) Pairwise temporal stability calculated by correlations of pairwise correlations across experimental states. Dark colored line represent mean correlation, shades represent 95th percentile. We tested pairwise stability of grid cell in the familiar environment prior to any change in environmental condition by identifying grid cells recorded in the familiar environment at any day prior to the day of the novel environment experiment. We only included cells that could be followed through the novel environment recording sessions. This is seen in the graph as Familiar vs Familiar I **d**). Distributions of Pearson correlation coefficients of pairwise correlations shows a significant reduction in correlation for the chABC treated group from the familiar to the novel environment (mean ± s.e.m.; Familiar I vs Novel I; Control 0.39±0.05(n = 51); chABC 0.26±0.04(n = 45), *U* = 1421.00, *p* = 0.045). Furthermore, grid cells from chABC treated rats show reduced correlations in the familiar environment after the animals have explored the novel environment (mean ± s.e.m.; Familiar I vs Familiar II; Control 0.43±0.05(n = 51); chABC 0.27±0.06(n = 29), *U* = 950.00, *p* = 0.036). Dark colored line show median. **e**) Pairwise spatial stability calculated across states as in **c**. **f**) Pairwise spatial correlations are reduced in chABC treated rats when going from the familiar to the novel environment. (mean ± s.e.m.; Familiar I vs Novel I; Control 0.28±0.05(n = 51); chABC 0.03±0.03(n = 45), *U* = 1495.00, *p* = 0.011). All tests are Mann-Whitney U tests. Spatial pairwise correlations are also reduced in the chABC throughout all sessions in the novel environment (Fig. S6).

### Reducing excitatory to inhibitory synaptic strength mimics the effect of PNN removal in a simulated grid cell network

Several mechanisms may underlie the reduced inhibitory firing rates observed after PNN removal, e.g. increased membrane capacitance (*29*), reduced excitability (*28*), increased diffusion of AMPA receptors (*27*). We observed structural changes in synaptic input to PV^+^ cells (Fig. 1d), and therefore wanted to test if changing synaptic weights in a continuous attractor network produced similar changes to grid cell spatial properties as we observed in experiments. We simulated populations of excitatory and inhibitory neurons in a continuous attractor model (*45*), where excitatory grid cells were connected to distant inhibitory neurons that inhibited excitatory neurons close by (Fig. 6a). Within this network, excitatory neurons only communicate through inhibitory neurons. Thus, to generate activity, the excitatory neurons received a non spatial external excitatory drive from a Poisson spiking generator (Fig. 6b). Because of the competitive inhibitory interaction between the excitatory cells, the activity of the network rapidly settles into a hexagonal grid pattern on the two-dimensional neuronal sheet (Fig. 6c). As expected, reducing excitatory to inhibitory synaptic strength produced a strong reduction in inhibitory firing rates, but it also led to reduced excitatory rates, similar to what we observed in the broad-spiking population (Fig. 6e, Fig. 1g). Grid cells showed increased out-of field rate and thereby lower specificity than control (Fig. 6d). The increased out-of field rate is caused by the reduction in firing rate of the inhibitory units in the simulation (Fig. 6e). Contrary to experimental data, we found increased peak rates of grid fields in the reduced-inhibition model (Fig. 6c). In a separate experiment we created a model where we increased the capacitance of inhibitory units aiming to match findings from an in-vitro study where PNNs were removed ((*29*)) (data not shown). Interestingly, in the increased-capacitance model, peak rates were reduced in grid fields similar to what we find in experimental data. All other results from the increased-capacitance model were similar to the results presented in Fig. 6, although less pronounced. A combination of effects on inhibitory neurons, i.e. reduced synaptic input and change in intrinsic excitability is likely to cause the observed effects in grid cell spiking properties in experimental data.

**Figure 6:**
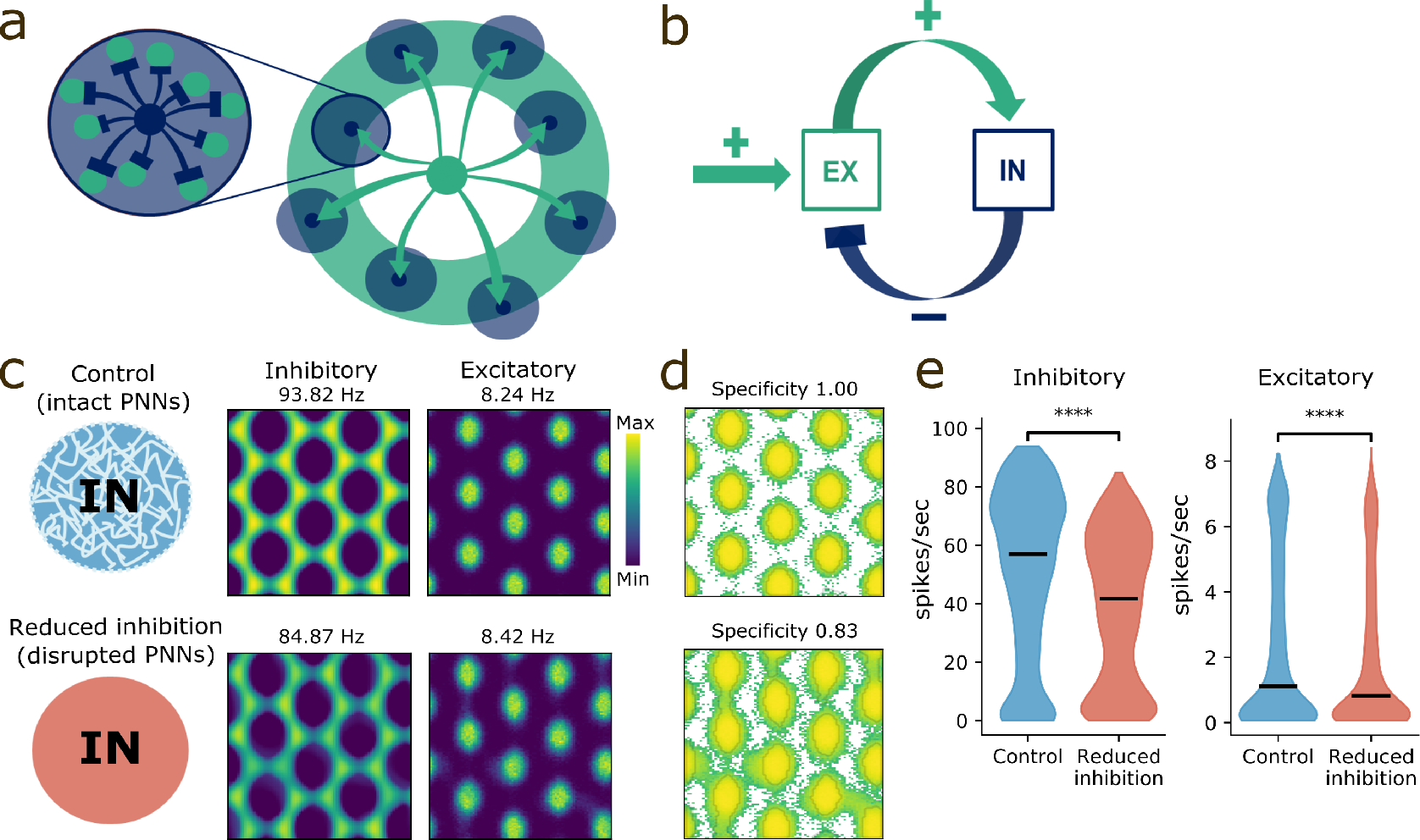
Continuous attractor model of the grid cell network with altered inhibition resembles experimental removal of PNNs. **a**) Schematic outline of the model. Each excitatory unit (green) is connected to distant inhibitory neurons (dark blue), which in turn inhibit excitatory neurons locally. **b**) A population of excitatory neurons (EX) receive external excitatory drive and feedback inhibition by a population of inhibitory neurons (IN). **c**) PNN removal was simulated by reducing the excitatory to inhibitory synaptic strenght (lower panel). Rate maps of excitatory neurons show grid pattern for both control- and the reduced inhibition network. **d**) The grid cells in the reduced inhibition network show increased out-of field rate and thereby lower specificity than control, as seen with logarithmic color scale. Note the increased number of green spikes representing rate outside the field. **e**) Average firing rate of both inhibitory and excitatory neurons are lower in the reduced inhibition network, (mean ± s.e.m., inhibitory control; 50.49±0.35, inhibitory reduced inhibition; 37.82±0.30*U* = 29940515*n* = 6858 and excitatory control; 2.37 ± 0.03 excitatory reduced inhibition; 1.95 ± 0.03*U* = 15509839.5*n* = 5398, *p* < 0.0001, Mann-Whitney U test).

### PNN removal in MEC changes the hippocampal place code

Grid cells are generally assumed to be the primary determinant of place cell firing (*5, 8, 13*), although this notion has been challenged in both experimental and computational work (*10, 46, 47*).

To test if the observed changes in grid cell activity after PNN removal affected the hippocampal place code, we recorded 185 units from hippocampus area CA1 (10 animals of which 4 had been treated with chABC in MEC). In the familiar arena we found that spatial specificity was significantly reduced in animals with disrupted PNNs due to a combination of reduced in-field rate and increased out-of-field rate (Fig. 7b, Table S2). These results are similar to what we observed for grid cells in MEC, only changes in in-field and out-of-field rates were more pronounced in place cells.

**Figure 7:**
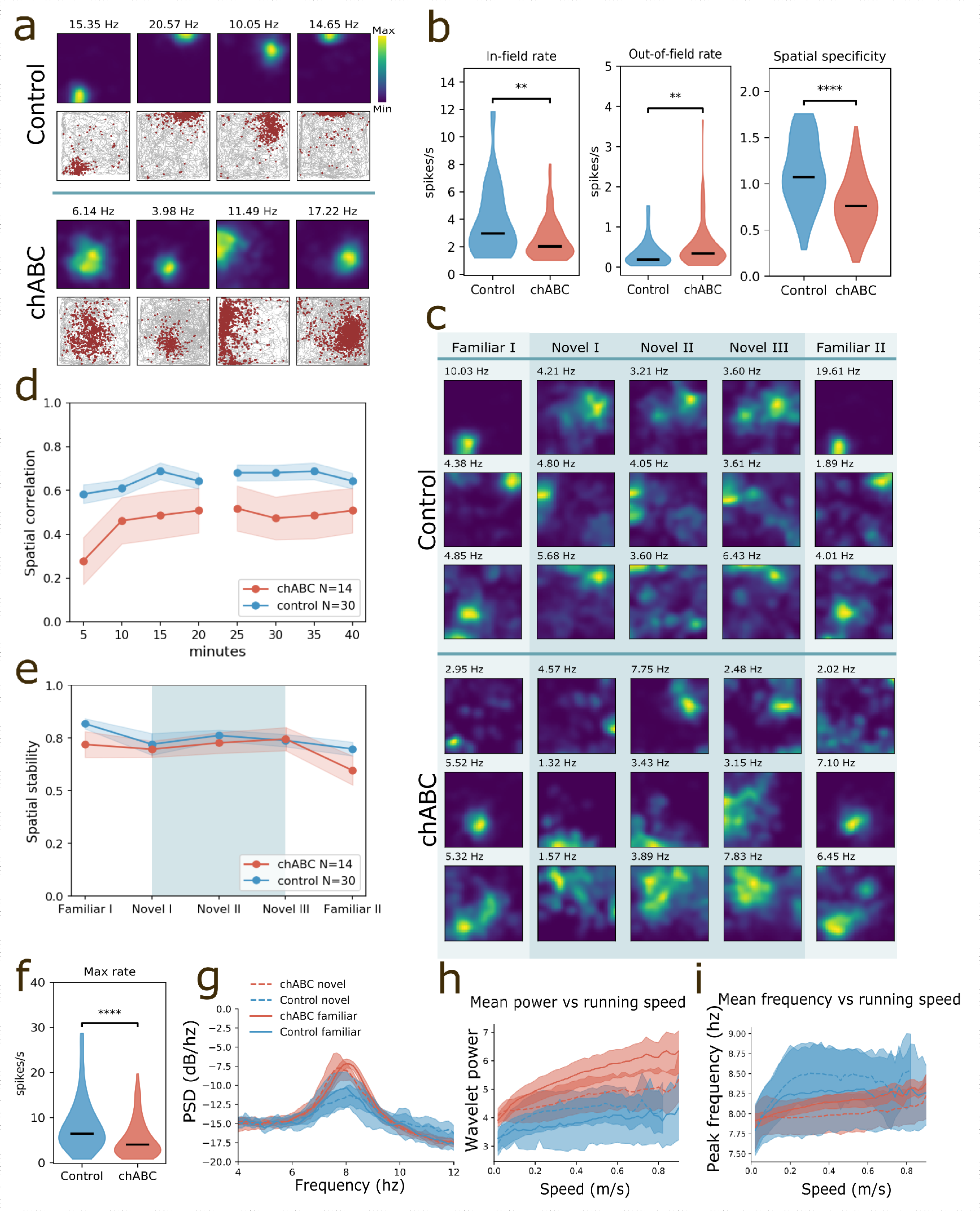
Altered hippocampal place code after PNN degradation in MEC. **a**) Examples of four place cells recorded in control and chABC treated rats. Top row shows color coded rate maps and bottom row shows the animals’ trajectory with spikes superimposed. Maximum spike rate is denotet above the rate map. **b**) Place cells show reduced in-field rate (mean ± s.e.m.; Control 3.78±0.12; chABC 2.54 ± 0.15, *p* = 0.003) and increased out-of field rate in chABC treated rats (mean ± s.e.m.; Control 0.34 ± 0.05; chABC 0.50 ± 0.05, *p* = 0.004, Mann-Whitney U test). **c**) Rate maps of five consecutive 20 minutes recording sessions from six individual place cells, recorded in control animals (top) and chABC animals (bottom) during a novel environment experiment. Rows correspond to individual units and columns to environment condition. **d**) Spatial correlation between blocks of 5 min recordings in a novel environment (Novel I & Novel II) against the last 20 minutes (Novel III), (mean ± s.e.m.; Control 0.65 ± 0.03; chABC 0.46 ± 0.09, main effect of group: *F*(1, 40) = 6.011, *p* = 0.0187, Control *n* = 30; chABC *n* = 14, two-way repeated measures ANOVA with group and time as factors). **e**) Within a session the place code is stable for both groups measured as spatial correlation between first and last 10 minutes of the session. **f**) Maximum spiking rate is strongly reduced in the novel environment in animals treated with chABC (mean ± s.e.m.; Control 16.38±3.02; chABC 5.30±0.48, *p* < 0.001, Mann-Whitney U test). **g**) Spectrogram showing that the power of theta oscillations are stronger in chABC treated animals. **h**) Mean theta power in hippocampus correlated with running speeds from familiar and novel environment recordings. **i**) Peak theta frequency increases with running speed. Violin plots show min to max and median (black line). Width of graph corresponds to number of samples for each value. \**p* < 0.05, \*\**p* < 0.01, \*\*\**p* < 0.001, **** *p* < 0.0001.

Next, we introduced the animals to a novel environment to test how place cells responded to decreased stability in MEC. Both control and chABC animals displayed hippocampal remapping. The place cells in the chABC treated group displayed reduced spatial correlations throughout the entire novel environment recording session (Fig. 7c, d), although individual units were highly variable. Despite reduced spatial correlations, the spatial stability within sessions was almost identical between the groups (Fig. 7e). Notably, we found greatly reduced maximum firing rates in chABC animals in the novel environment compared to controls (mean ± s.e.m.;control 16.38 ± 3.02, n = 67; chABC 5.30 ± 0.48, n = 77, *U* = 3916, *p* < 0.001, Mann-Whitney U test)(Fig. 7f). Maximum firing rates of place cells is normally expected to increase in the novel environment (*46, 48*), which indicates that removing PNNs affected the input from MEC sufficienty to impair rate change in hippocampal place cells.

Although PNNs were only removed in MEC, the LFP recorded in hippocampus also showed increased power at the peak of theta (Fig. 7g). Similar to MEC, theta power was increased for all running speeds, but only in the familiar environment (Fig. 7h, Fig. S5d). The peak frequency of the theta oscillations increased strongly when the animals’ behavior shifted from sessile to movement, but quickly reached a plateau for the control animals, while the injected animals showed a more linear correlation between theta frequency and running speed (Fig. 7i). Interestingly, the correlation between frequency and running speed (frequency score) was decreased in chABC treated animals in hippocampus, which is opposite from what we see in MEC (Fig. S5c).

## Discussion

The grid cell network in MEC has been under intense investigation since its discovery (*6*), but little is known about how grid cells achieve their extraordinary stability across time and environments. Perineuronal nets have been shown to stabilize synapses and limit plasticity in several brain areas. Here we investigated how the grid cell network is affected by enzymatic degradation of PNNs. We show that degradation of PNNs changes grid cell network dynamics by altering the temporal relationship between grid cells and impairing representations of novel environments. Results from simulating the grid cell network in a continuous attractor model indicate that reducing the level of inhibition is sufficient to produce impairments in grid cell spatial specificity, similar to what we observe in experimental data after removal of PNNs. These results support the notion that PNNs stabilize the network through its effects on inhibitory, putative PV^+^ neurons, and suggest a prominent role for PNNs in maintaining the structural and functional organization that shapes the activity of microcircuits in MEC.

### Reduced inhibition and altered connectivity weakens grid cell precision

Removal of PNNs in MEC caused both structural and functional changes in the grid cell network. The number of inhibitory VGAT-positive puncta on PV^+^ cell somas decreased, but it did not result in an increased firing rate of narrow-spiking putative inhibitory neurons. On the contrary, when recording from neurons in MEC we found that removal of PNNs caused a reduction in the activity of inhibitory neurons, similar to what has been found in other brain areas (*24, 49*). This affected the spiking and spatial precision of grid fields in the familiar environment. However, we did not observe impairment in spatial stability or lower gridness scores in the familiar environment that resemble the instability seen in juvenile grid representations (*9, 11*). This indicates that the connectivity supporting the spatial spiking patterns of grid cells remained stable enough to maintain precise spatial maps as long as the environment was familiar.

### PNNs are necessary for grid cell representations of novel environments

The most pronounced effects of PNN removal were observed for representations developed in a novel environment (Fig. 3). Without PNNs, grid cells failed to establish stable representations of the novel environment. In addition, the novel environment experience interfered with representations of the well-known arena (familiar environment). The latter resembles previous work from the amygdala where extinction training in adult animals after fear conditioning, combined with PNN removal, completely replaces the original memory trace instead of creating a separate extinction representation (*50*). Together with our data, this suggests that intact PNNs facilitate storage and maintenance of separate memory traces so that newly formed memories do not interfere with previously stored experiences. For the grid cell network, this appears as the ability of the network to maintain representations of a seemingly infinite number of different arenas as well as other non spatial modalities (*51*).

### Loss of PNNs in MEC destabilizes hippocampal place cells

While place cells can form even if input from MEC is minimized, they display reduced spatial precision and stability under these circumstances (*8*). Specifically increasing the scale of grid cells or impairing grid cell spiking patterns reduce the long-term stability of place cells (*46, 52*), making it evident that the properties of grid cell input plays an important role for place cell spatial coding. Our data showed that the manipulations in MEC was reflected in recorded place cells, thus giving support to this assumption.

In contrast to the highly organized representations in MEC, the hippocampus represents different environments by recruiting different subpopulations of place cells (*53, 54*). It is likely that high levels of plasticity in the hippocampal network enables place cells to rapidly reorganize and produce a vast number of orthogonal representations. Perhaps surprisingly, place cells can produce new and distinct spatial representations when the firing pattern of grid cells are degenerated, although with reduced stability and reduced rate change in the novel environment (*46*). When the chABC treated rats were introduced to the novel environment we observed a reduction of spatial correlations in the place cell population similar to what we observed in grid cells. In addition, we found a striking lack of change in maximum firing rate. This suggests that even when grid cells produce grid cell spiking patterns, the spatial stability and temporal precision of grid cells affects place cell coding in novel environments under normal conditions. Taken together, our data from recordings in hippocampus when PNNs were removed in MEC indicate that place cells rely on adequate grid cell activity for producing precise spatial representations in both familiar and novel environments.

### Stronger theta oscillations after PNN removal

We show that PNN removal leads to reduced bursting and spiking variability in grid cells. This in itself could give rise to more synchronous firing patterns leading to the increased theta power we see in animals treated with chABC. Since increased theta power is associated with enhanced plasticity and increased synchronization of interconnected input (*55, 56*), we find it likely that the high theta power in combination with reduced inhibitory activity reflects a state of increased plasticity in the MEC network.

Another possibility is that the increased theta power we observed after PNN removal could stem from a reduction in the number of inhibitory (VGAT) synapses from the medial septum to PV^+^ cells in MEC. However, this would likely reduce theta power due to reduced disinihibition and is thus an improbable explanation for the observed changes in theta.

It has been suggested that environmental novelty is signaled by a sharp reduction of the hippocampal theta frequency, which gradually disappear with repeated exposures to the environment (*57*). In line with this we observed a reduction in frequency upon transition from the familiar environment to the novel environment (Novel I), for both control and chABC treated animals (Fig. 4). In the subsequent recordings in the novel environment (Novel II and III) the peak theta frequency increased as the environment became more familiar. Interestingly, the increase was faster and more pronounced in the control animals. Furthermore, the control animals returned to their baseline frequency when returning to the familiar environment (Familiar II), while the theta frequency continued to increase in most chABC treated animals. Together with the single unit recordings, this supports that exposure to the novel environment disturbs the representations of the familiar environment in animals lacking PNNs in MEC.

### Reduced spatiotemporal spiking relationship in grid cells after removal of PNNs

We found that spatiotemporal correlations between pairs of grid cells are maintained in control animals when they are introduced to a novel environment, but not in animals where the PNNs were removed (Fig. 5). To our knowledge, this is the first manipulation that alters the spatiotemporal relationship between grid cells across environments. Recently, grid cells were shown to exhibit a temporal correlation structure that transcends across behavior states (*58, 59*). In addition, the relationship between grid cell pairs remains highly stable across environments despite huge changes in single cell responses (*6, 16*). This indicates that the local circuitry supporting grid cell firing normally exhibits very little plasticity, possibly due to the presence of PNNs.

Interestingly, we also find that pairwise spatial correlations are reduced in chABC treated animals for all sessions in the novel environment (Fig. S6). This provides further support for the observations of reduced stability of newly formed grid fields after PNN removal. The reduced pairwise correlations could both be a result of lower inhibition and changes in synaptic connections onto inhibitory neurons. Either possibility is likely to reflect an increase in network plasticity.

### Continuous attractor hypothesis

Computational models have shown that inhibitory connections are sufficient to generate the hexagonal structure of the grid pattern (*17, 35, 60*). Thus, the dispersed grid cell patterns we observed after PNN removal is likely a consequence of reduced inhibition, especially since the increased out-of-field spiking occurs where grid cells presumably rely on inhibitory domination in normal circumstances. This result is consistent with a previous report showing that pharmacogenetic silencing of PV^+^ interneurons selectively affects grid cells in a similar manner(*12*). However, in contrast to our findings, this study saw an increase in mean spike rate of grid cells, but no change to the maximum spike rate. This indicates that removal of PNNs causes alterations in network properties that can not be explained solely by an acute reduction in PV^+^ cell activity.

In line with experimental data, we observed that lowering excitatory synaptic strength onto inhibitory neurons caused reduced inhibitory spiking in the continuous attractor model (Fig. 6c). Furthermore, it also led to a decrease in mean spike rate of excitatory neurons, similar to what we observed in the experimental data. This result is, at first glance, counter intuitive as reduced inhibition would be expected to increase excitatory activity. However, the dynamical interactions in the model and the spatial outreach of connections have a large influence on activity levels of both inhibitory and excitatory neuron populations (data not shown).

Our results from the familiar environment experiments could largely be explained by reduced firing rates in both inhibitory and excitatory neurons, as shown by the in-silico data. However, in a novel environment, increased plasticity seems to be relevant because both reduced spatial correlation and reduced gridness is not observed until the animals are introduced to a novel environment. In addition, disrupted stability in cross correlation structure is clearly seen when there is an abrupt change in the environment, i.e. when animals go from Familiar I to Novel I and later when they are reintroduced to the familiar environment (Familiar II).

Accumulating evidence points to a low dimensional continuous attractor to be present in the grid cell network (*16, 58*). In this theory the co-activity of pairs of neurons remains stable throughout experiments with varying environmental conditions, meaning that the population activity resides in a low dimensional manifold even if single neuron activity changes dramatically. Our results indicate that this manifold is partially disrupted through destabilization of pairwise correlations, which again suggest that the attractor dynamics of the population is shifting away from its stable state. It is not fully clear, however, why this only happens when the animals are introduced to a novel arena. One possibility could be that novel environment exploration stimulate synaptic plasticity that is normally restricted in adult animals. Nonetheless, removal of PNNs could be removing the brakes on plasticity, allowing attractor dynamics to shift away from the stable state once the animals are introduced to the novel environment.

Computational models have shown that the spatial firing pattern of grid cells can develop and be supported by external input from place cells (*47, 61*). However, in order for grid cells to share alignment as shown in numerous experimental studies, it seems likely that some form of attractor dynamics is necessary (*62*). The seemingly hard-wired nature of the MEC L2/3 network could therefore be necessary to maintain the relative relationship between grid cells.

By removing PNNs, we show that the spatial representations of individual grid cells remain relatively stable, while pairwise spatiotemporal relationships are decreased in strength. Hence, these data provide support for a local, low-plasticity attractor network and the role of inhibitory control in grid cell firing postulated by continuous attractor models (*17, 35, 60*).

In summary, we show here that PNNs are essential to stabilize the grid cell network and to ensure proper function when the network is challenged to create representations of novel environments. By stabilizing connections and supporting proper inhibition, intact PNNs may be essential to efficient navigation.

## Methods

### Subjects

This study used recording data from 23 male Long–Evans rats (3–8 months old, 350–550g at surgery). After surgeries the animals were housed individually in transparent Plexiglas cages (45 × 30 × 35 cm) in a temperature- and humidity-controlled vivarium. All rats were maintained on a 12-h light/12-h dark schedule. Testing occurred in the dark phase. The rats were kept at 85–90% of free-feeding body weight and food deprived 18–24 h before each training and recording trial. Water was available ad libitum. Experiments were performed in accordance with the Norwegian Animal Welfare Act and the European Convention for the Protection of Vertebrate Animals used for Experimental and Other Scientific Purposes.

### Surgical procedures

All surgical procedures were performed in an aseptic environment. Rats were anesthetized with isoflurane mixed with air (5% induction, 1.5-2% for maintenance) and immobilized in a stereotaxic frame (World Precision Instruments Ltd, Hertfordshire, UK). They were given subcutaneous injections of buprenorphine (0.04 mg/kg) and local subcutaneous injections of bupivacaine/adrenaline (Marcain adrenaline, 13.2 mg/kg) in the scalp before surgery began. The scalp was shaved and cleaned with ethanol and chlorhexidine. Heart rate and core temperature were continuously monitored throughout the operation through a MouseStat system (Kent Scientific, CT, USA), the latter in a feedback mechanism to a heating pad. In addition, the hind paw withdrawal reflex was used to assess the depth of anesthesia.

#### Microinjection of Chondroitinase ABC (chABC)

Craniotomies were made bilaterally above the MEC, using a hand-held Perfecta-300 dental drill (W & H Nordic, Täby, Sweden). Injections were made in concomitance with tetrode implantation. Protease-free Chondroitinase ABC (chABC) from Proteus vulgaris was purchased from Amsbio (Abingdon, UK) and diluted in filtered 1X phosphate-buffered saline (PBS) to a concentration of 0.05 U per μl (Pizzorusso et al. 2002). Glass pipettes with a 15-20 μm opening diameter were backfilled with mineral oil, assembled into a NanoJect II microinjector (Drummond Scientific Company, PA, USA) and loaded with chABC. Injections were done in 2 depths at each location (2500 and 3200 μm below the dura). The locations were at AP 0.5 anterior of the transverse sinus, and ML 4.4 and 4.7 mm relative to the midline. A total volume of 3.1 μL was injected in each hemisphere. The injections were made step-wise over two minutes at each position, and the pipette left for another two minutes before retraction.

#### Electrode implantation

Tetrodes were implanted above MEC at AP 0.4 ± 0.1 mm in front of the transverse sinus and ML 4.5 ± 0.1 mm relative to the midline. Tetrodes implanted above hippocampus were placed at AP Bregma −3.8 ± 0.2 mm and ML 3.1 ± 0.1 mm. The depth of implantation was 1800 μm measured from the surface of dura. Jeweler’s screws fixed to the skull served as ground electrodes. The microdrives were secured to the skull using jeweler’s screws and dental cement. All animals were given a subcutaneous injection of carprofen (5 mg/kg) at the end of the surgery, and the edge of the wound was cleaned and local anesthetic ointment Lidocain was applied. This was repeated for three days after surgery.

### Extracellular recordings

Electrophysiological recordings were performed within 14 days after chABC injections since it has been shown that less than 40% of PNNs are reassembled by that time (*24*). The recording system used was daqUSB, provided by Axona (Herts, UK). Signals were amplified 8000-15000 times and band-pass filtered between 0.8 and 6.7 kHz. Triggered spikes were stored to disk at 48 kHz (50 samples per waveform, 8 bits/sample) with a 32-bit timestamp (clock rate at 96 kHz). Spike waveforms above a threshold of 50 μV were time-stamped and digitized at 32 kHz for 1 ms, and saved to the hard-drive for offline analysis. One channel in each hemisphere served as a reference electrode to record LFP; LFP signals were amplified 3000 times, low-pass filtered at 500 Hz and stored at 4.8 kHz (16 bits/sample).

#### Recording procedure

Prior to surgery the animals were habituated to a recording environment that during the experiments became their familiar environment. The environment consisted of a 1×1 meter black box containing a white A4 sheet on one of the walls serving as a local cue. Animals were motivated to explore the environment by chocolate crumbles that were thrown at random intervals into the recording arena. During recording sessions, animals ran in the arena for sessions lasting from 10 to 20 minutes (depending on how fast they could cover the entire environment). Between sessions, the tetrodes were adjusted downwards in steps of 50Um and the animals allowed to rest in their home cage for 20-30 minutes.

#### Novel environment recordings

When the tetrodes were located in MEC (usually when three or more grid cells could be recorded simultaneously), novel environment experiments were conducted. The novel environment consisted of an identical recording setup as the one in the familiar environment but located in a different experiment room. In addition, the cue card was placed on a different wall in the recording arena relative to the familiar environment. This paradigm is shown to cause consistent global remapping of spatial representations in both hippocampus and MEC (*15, 53*).

First, a recording session of 20 min was conducted in the familiar environment followed by 3×20 minutes recording sessions in a novel environment. Lastly, the animals were put back in the familiar environment for an additional 20 min session. The animals rested for 5-10 minutes in the homecage between sessions. This experimental procedure was repeated for three consecutive days, with the exception that the novel environment recording on day 2 and 3 consisted of only one 20 minute long recording session.

#### Spike sorting

Single units were isolated using the offline manual spike sorting software Tint (Axona). Clusters were manually isolated using 2D projections the multidimentional parameter space (waveform amplitudes). Cross-correlation and autocorrelation were used as additional tools for assessing the quality of separation. Spikes falling within the 2-3 ms refractory period of neurons were considered to belong to other units and if further spike sorting did not remove these, the unit was discarded. Identification of the same units in successive recording sessions (novel environment experiments) was done by manual inspection of cluster location and waveform, in addition to comparing the behavioral outputs of single units.

### Histology and identification of recording position

At the end of the experiments, animals were deeply anesthetized by an intraperitoneal injection of pentobarbital sodium (50 mg/kg) and intracardially perfused with 0.9% NaCl followed by 4% paraformaldehyde (PFA) in 1X PBS. The brains were dissected out and post-fixed in 4% PFA overnight. They were cryoprotected in 30% sucrose in 1X PBS for three days, and 40 μm sagittal (MEC) or coronal (hippocampus) sections were cut with a cryostat. Staining procedures were performed on free-floating sections under constant agitation unless mentioned otherwise. The lectin *Wisteria floribunda* agglutinin (WFA) was used to visualize PNNs.

The sections were rinsed times in 1X PBS and blocked with 2% goat serum with 0.3% Triton X-100 in 1X PBS for 1 hour at room temperature. The sections were then incubated with biotin-conjugated WFA (#L1516, Sigma-Aldrich Chemie, 1:200)in blocking solution overnight at 4°C. On the following day, the sections were rinsed with 0.3% Triton X-100 in 1X PBS, endogenous peroxidase activity was quenched for 3 minutes with 2% H202 in dH2o, and the sections incubated with ABC solution (ABC Peroxidase Standard Staining Kit, Thermo Fisher Scientific). After rinsing the sections in Tris nonsaline (TNS), staining was visualized by adding a 3,3-diamonobenzidine hydrochloride (Sigma-Aldrich Chemie) solution (0.67 mg/ml 0.05M Tris-HCl, with 0.8μl H202/ml) for 3-10 minutes. Sections were rinsed in TNS, mounted on coverslips and dried for 1 hour. They were then left overnight in a solution of 1:1 choloroform and 96 % ethanol. The following day, the sections were rinsed in dH2O and counterstained for Nissl substance using Cresyl Violet and coverslipped with Entellan. Tetrode tracks were identified, measured and photographed through an Axioplan 2 microscope (Carl Zeiss, Oberkochen, Germany). High-resolution images were then stitched together using the MosaiX extension in the AxioVision software (Carl Zeiss, Oberkochen, Germany).

#### Immunohistochemistry

A subset of sections from each animal were used for visualizing the chABC treated area in MEC. Sections were prepared as described above. After blocking, the sections were incubated with WFA (#L1516, Sigma-Aldrich, 1:200) and chondroitin-6 sulfated stubs (MAB 2035, Millipore, 1:1000) in blocking solution overnight at 4°C. The next day the sections were rinsed with 0.3% Triton X-100 in 1X PBS and incubated for two hours with Streptavidin Alexa 488 (#S-11223, Life, 1:1000) and donkey-anti mouse Alexa 594 (#A-21203, Life, 1:1000) in 1X PBS. Sections were rinsed in 1x PBS, mounted to Superfrost Plus slides, washed in dH20 and coverslipped with FluorSave Reagent (#345789, Millipore).

#### Analysis of synaptic boutons

Animals were injected with chABC in MEC of one hemisphere and artificial cerebrospinal fluid (aCSF) in one hemisphere. After surgery, animals were kept in their home cage for five days until they were transcardially perfused and the brains were sectioned as described above. Prior to antibody incubation, sections were blocked with 2% BSA and 0.3% Triton x-100 in 1x PBS for one hour.

Two sections from each hemisphere representing the medial-center and lateral-center parts of the injection site (approximately 100-120 μm apart), were incubated overnight with goat anti-PV (#PVG-214, Swant, 1:2000), biotin conjugated Wisteria floribunda agglutinin (#L1516, Sigma-Aldrich, 1:200) and either rabbit-anti vGlut1, rabbit anti-vGlut2 or rabbit anti-vGat (1:5000, synaptic vesicle antibodies kindly donated by Dr. Farrukh Chaudry, UiO) in blocking solution containing 0.2% sodium azide. The next day, sections were rinsed with 1x PBS and incubated with donkey anti-goat Alexa 594 (#A-11058, Life, 1:1000), Streptavidin Alexa 647 (#S-21374, Life, 1:1000) and chicken anti-rabbit Alexa 488 (#A-21441, Life, 1:4000) in 1x PBS for two hours. Sections were rinsed in 1x PBS, mounted to Superfrost Plus slides, washed in dH20 and coverslipped with Fluoromount mounting medium containing DAPI (#0100-20, AH Diagnostics).

Images were acquired with an Andor Dragonfly spinning disc microscope using the Fusion software, with a Zyla 5.5 sCMOS camera covering 2048 × 2048 pixels. Overview images of entire slides containing two sections per antibody combination from one hemisphere of one animal, were acquired using a 4X objective (NA 0.2). This verified that chABC had degraded all PNNs in mEC in all animals, and verified the needle tracks from aCSF injections in sham treated tissue.

Landmarks from these images were used to verify that all high magnification images were acquired from within the mEC. Detailed images of PV^+^ somas and markers of synaptic vesicular transporters were acquired through a 60x objective (1.4 NA). Z-stacks with a step size of 0.16 yU,m were acquired through the entire cell soma. One to three somas were included in each stack.

Images were analyzed in a similar fashion to (*63*) using Imaris 9.1.2. First, regions of interest containing each soma were selected. Second, synaptic boutons were detected using the built-in spot detection algorithm with background subtraction. Finally, boutons in contact with each respective soma were labeled using the Spots close to Surface-tool, with maximum distance from center of the spot to the surface set at 0.8 μm. Each set of spots and soma were manually inspected and revised after processing. Data from all animals in each group (chABC or aCSF) were pooled prior to statistical analysis.

### Quantification and statistical analysis

All tests are two-sided. When testing differences in means between the experimental groups, we tested for normality in a subset of data using the Shapiro-Wilk test in Sigmaplot. For the subset of data we tested, normality failed in all, hence we only used non-parametric tests (Mann-Whitney U test) for all data. To test if grid cell parameters were consistent across animals within a group and that results were not caused by random variations, we performed a permutation resampling test in addition to Mann-Whitney U test. A low *p* value allowed us to reject the hypothesis that data could be randomly drawn from either group. For testing spatial correlations over time in the novel environment, we performed ANOVA with repeated measures and Šidák’s multiple comparisons post hoc test, using GraphPad Prism 8. All other analyses were performed in Python.

#### Spiking rate maps

To produce Spiking rate maps, we divided the arena into equally sized bins. For each bin, we counted the number of spikes and the time spent within the bin to produce a spike map and an occupancy map, respectively. The bin size was set to 2 cm × 2 cm. We further smoothed the spike map and the occupancy map individually by using the convolution of a two-dimensional Gaussian kernel. By dividing the value of each bin in the smoothed spike map by the corresponding bin in the smoothed occupancy map, we produced a smoothed rate map in two versions, one with standard deviation *σ* = 3 cm and one with standard deviation *σ* = 5 cm.

#### Autocorrelogram

To create a spatial autocorrelogram used for visualization and subsequent analyses, we correlated the smoothed rate map with itself using the `scipy.signal.fftconvolve` function from the SciPy package with the mode parameter set to mode = ‘full’.

#### Gridness score

To calculate the gridness score of each rate map, we first identified the central peak and the six closest peaks in the autocorrelogram by finding the maxima of the autocorrelogram and calculating the distance to the center of the autocorrelogram to each maximum. The center peak was selected as the peak closest to the center of the autocorrelogram. We then masked out the center peak with a disk centered on the center peak with a radius of half the distance to the closest peak. To mask the area of the autocorrelogram outside a circle going around the outside of the outermost of the six closest peaks, we masked everything outside a disk centered on the center peak with a radius 3/2 times the distance to the outermost of the six closest peaks. Next, the masked autocorrelogram was rotated by increments of 30° up to 150°. For each rotation, we calculated the Pearson product moment correlation coefficient against the originally masked autocorrelogram. Finally, the gridness score was calculated by taking the lowest coefficient found with rotations 60° and 120° and subtracting the highest coefficient found with rotations 30°, 90° and 150°.

#### Shuffling

To obtain null distributions such that statistical properties, such as gridness, could be measured against, we generated a randomized spike train for each registration using the function `spike_train_surrogates.dither_spike_train` from the Elephant electrophysiology analysis toolkit.^1^ We generated *n* = 1000 randomized spike trains for each session using a shift of 30 s and with the edges keyword set to `edges=True`.

#### Spatial correlation

Spatial correlation across trials was calculated according to the procedure for calculating intertrial stability in (*37*). To calculate the reliability of spatial firing between entire or partial trial against a reference, we calculated the Pearson product moment correlation coefficient of the rate map from the trial against the rate map of the reference using the corrcoef function from the NumPy Python package.

#### Spatial stability

Spatial stability was calculated as within-trial spatial correlation according to the procedure for calculating intra-trial stability in (*37*). To calculate the reliability of spatial firing within a trial, we calculated the Pearson product moment correlation (see inter-trial stability above) of the first 10 minutes of the trial against the last 10 minutes of the trial.

#### Spatiotemporal pairwise cross correlations

First only grid cell pairs were selected from the same recording and hemisphere. Temporal pairwise cross correlations were calculated by first calculate the instantaneuous firing rate by convolving the spike times with a Gaussian kernel 10 ms width using `statistics.instantaneuous_rate`^2^ at temporal differences from −1 to 1 seconds using `numpy.correlate` with `mode=‘full’` and `method = ‘fft’`. The rates were zscored and one of the two zscored rates were scaled by it’s size to acheive a crosscorrelation value between −1 and 1. Then, the Pearson correlation coefficient of pairwise cross correlations was used to quantify the similarity across experimental states using `numpy.corrcoef`.

To assess the spatial cross correlations we first calculated the rate maps, then for each pair we calculated the 2D cross correlation function using `astropy.fft_convolve`^3^ with `boundary=‘wrap’, normalize_kernel=False, nan_treatment=‘fill’`. Each rate map were zscored and one of the two zscored rates were scaled by it’s size to acheive a crosscorrelation value between −1 and 1. Then, the Pearson correlation coefficient of pairwise cross correlations (reshaped to 1D) was used to quantify the similarity across experimental states using `numpy.corrcoef`.

To control for possible effects due to changed theta acitivity in the experimental group a Butterworth bandstop filter was used to filter out frequencies between 4 and 10 Hz, this did not affect results significantly. Furthermore, to control for effects due to differences in out of field spikes between the experimental and control group we also removed all spikes landing outside fields before calculating rates, this did also not change results significantly.

#### Identification of firing fields

To estimate the firing within and outside fields, we first identified the individual fields in the rate map. Following the protocol of (*64*), we first identified a global field radius as 0.7 times half the distance from the center peak to the closest peak in the autocorrelogram. Next, we identified all the peaks in the rate map before excluding the lowest of any two peaks within a distance shorter than the global field radius.

To define the extent of each field, we calculated the Laplacian, ∇^2^, of the smoothed rate map to obtain its curvature and excluded regions with a positive Laplacian, which are the valleys of the rate map. The Laplacian was calculated using the `ndimage.laplace` function from the SciPy Python package.

To separate and label the remaining regions, we used the ndimage.label function. Any regions with an area less than 9 bins were excluded. The areas were then sorted based on the mean firing rate in each area.

Fields were defined to be any labeled region found after taking the Laplacian that corresponded with a non-excluded peak from the protocol of (*64*). The fields were used in subsequent analyses to identify in- and out-field spikes.

#### Spatial specificity

Spatial specificity is a measure of the firing ratio between in-field firing and out-field firing and is defined as

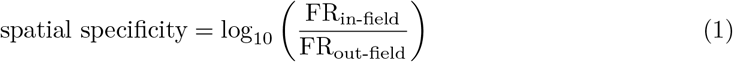

#### Spatial information

Spatial information is an estimate of to which degree the animal’s position can be predicted based on the firing of the cell and is given in bits per second. Spatial information was found by

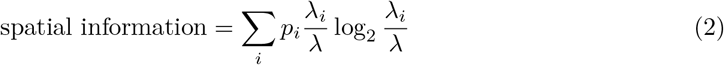

where *p_i_* is the probability of the animal being in bin *i*, given by the occupancy map divided by the total session time, *λ_i_* is the firing rate in bin *i*, and *λ* is the mean firing rate.

#### Coefficient of variation

The coefficient of variation (CV) for interspike intervals (ISI) was calculated as the standard deviation of the ISI distribution σ_ISI_ divided by the mean of the ISI distribution *μ*_ISI_:

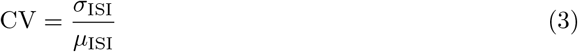

To reduce bias in ISI that arises from animals traveling between receptive fields, we calculated only in-field CV-values by extracting one CV-value for each pass through the fields. This was compared with two other methods. In the first, a CV-value was calculated for the entire session. The second followed the protocol of (*60*), where sections of the recording where the speed of the rat was less than 8cm/s were selected. A CV value was then calculated for the sections with a longer duration than 0.2 s and at least 2 spikes.

See supplementary Fig. S4 for comparison of the three methods.

#### Bursting ratio

The bursting ratio was estimated by taking the interspike intervals (ISI) and defining any spike with a pre-ISI of more than 10 ms and a post-ISI less than 10 ms as a the start of a bursting event. Any spike with a pre-ISI and post-ISI of more than 10 ms was identified as a single spike event. All other spikes were assumed to be part of a bursting event and were not counted. The bursting ratio is then given by the number of bursting events *n_B_* divided by the number of single-spike events *n_S_*:

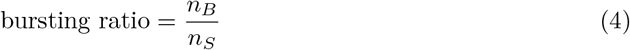

#### Waveform analysis

To separate between narrow spiking and broad spiking units we calculated the time from through to peak and time to cross the half width amplitude of the largest amplitude through from the mean waveform. In peak to through calculations, sampling period of each spike was increased 200 fold bu cubic interpolation to get an accurate measure of peak times. The half width crossing time was refined by a linear interpolation between crossings of the constant line of half amplitude. All interpolations were done with `scipy.interpolate.interpld.`

Finally we separated clusters with `scipy.cluster.vq.kmeans` separately on the control and chABC group.

#### LFP Spectrum analysis

To calculate the power spectrum versus running speed *v* we first calculated the instantaneous speed and interpolated linearly to match the sampling frequency of the LFP signal at 250 Hz. The power spectrum was calculated by means of a continuous wavelet transform on the z-scored LFP signal using a Morlet wavelet with nondimensional frequency *ω*_0_ = 80 (*65*) and the PyCWT library ^4^. Then a weighted histogram of of speed *v* with bin size 0.02*m/s* was calculated in the range *v* ∈ [0.02, 1]*m/s*. The weights were either the mean power of the wavelet spectrogram over frequencies *f* ∈ [4, 12] or the frequency at the maximum power withing the same frequency range.

The power spectral density (PSD) were calculated on z-scored LFP signal using the Welch method given by mlab.psd from the (*66*) package.

The confidence intervals presented in (Fig. 4, a-c) were calculated by bootstrapping each data point from each recording at the 95 % confidence level.

#### Continuous attractor model

Simulations were conducted to assess the effects of increased membrane capacitance due to removal of PNNs as reported by (*29*). To this end populations of excitatory and inhibitory neurons were simulated in a continuous attractor model. Neurons were modeled by the exponential integrate and fire model given as

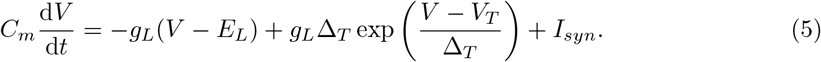

When the membrane potential *V* reaches the threshold *V_T_* there is an exponential increase in potential, with slope factor ∆_T_. The nonlinear spike is reset at *V_reset_* whereas the potential is reset to the equilibrium potential *E_L_*. Moreover, *C_m_* denotes the membrane capacitance, *g_L_* represents the membrane leak conductance.

The synaptic input current *I_syn_* is modeled by a difference of exponentials, also known as the *β* function given by

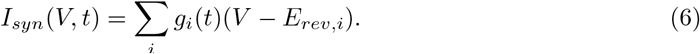

Here, *E_rev_* is the reversal potential, *i* denotes neuron number and the conductance *g* is given by

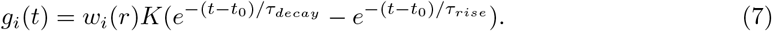

In Eq. (7), *ω*(*r*) is a spatial connectivity function, *t*_0_ represents onset time of a incoming spike, and *τ_decay_*, *τ_rise_* are the decay- and rise time-constants, respectively. Finally, *K* is a normalization factor, described in detail in (*67*)[Eq. 6.6].

To reach a continuous attractor state with hexagonal bump pattern, excitatory and inhibitory neurons were placed in grids of 100 × 100 each population forming a layer of extent 0 < *x* < 2*π*, 0 < *y* < 2*π*. Excitatory neurons were driven by a background Poisson noise process *p_rate_* and connected to the inhibitory layer through a doughnut shaped connectivity pattern with spatial connectivity function given by

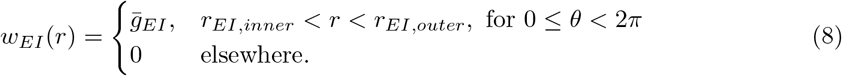

Here 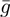 is the peak conductance amplitude, subscript *EI* indicates that the connectivity is directed from excitatory to inhibitory population. Position is given in polar coordinates with radial length 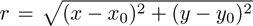 relative to neuron position *x_0_, y_0_* and angle *θ*. Inhibitory neurons were connected to excitatory neurons by

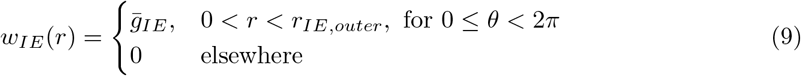

Topological connections were specified using the Topology package in NEST, and simulations were performed with NEST (*68*).

## Data and code availability

The source code used for analysis and raw data will be made available at the publication of the final manuscript.

## Acknowledgements

This work was funded by the Research Council of Norway (Grant No. 217920 to M.F, Grant No. 231248 to T.H) and the University of Oslo’s Strategic Research Initiative CINPLA. The authors wish to thank Anne Marthe S. Kvello for help with pilot experiments, and Jennifer Hazen and Benjamin A. Dunn for discussions and useful suggestions to the manuscript.

## Author contributions statement

A.C.C., M.F. and T.H conceived of and designed the project, A.C.C., K.K.L. and J.S.B. conducted the experiments, M.E.L performed the simulations, A.C.C, K.K.L., M.E.L., S-A.D., H.S. and T.H. analyzed the results. A.C.C., K.K.L., M.E.L., S.A.D., H.S, M.F: and T.H wrote the manuscript. All authors reviewed the manuscript.

## Supplementary information

This section contains Supplementary tables 1-2 and Supplementary Figure 1-10.

**Table S1:** Šidák’s multiple comparisons post hoc test for spatial correlations in a novel environment (mean diff and 95 % CI of diff).

| Šidák's multiple comparisons test |  |
| --- | --- |
| 0-5 min | 0.06 - 0.13 to 0.25, p = 0.975 |
| 5-10 min | 0.17 - 0.07 to 0.40, p = 0.343 |
| 10-15 min | 0.28 0.08 to 0.48, p = 0.002 |
| 15-20 min | 0.28 0.09 to 0.48, p = 0.002 |
| 20-25 min | 0.21 0.02 to 0.41, p = 0.030 |
| 25-30 min | 0.24 0.05 to 0.43, p = 0.008 |
| 30-35 min | 0.28 0.08 to 0.48, p = 0.003 |
| 35-40 min | 0.28 0.09 to 0.48, p = 0.002 |

**Table S2:** Firing properties of place cells in a familiar arena for control and chABC groups (mean ± s.e.m. (n), Mann-Whitney U (U, p) test and Permutation resampling (diff, p)).

|  | Control | chABC | MWU | PRS |
| --- | --- | --- | --- | --- |
| Average rate | 0.61 ± 0.07 (52) | 0.81 ± 0.07 (113) | 2454.00, 0.090 | 0.11, 0.213 |
| Max rate | 8.32 ± 0.88 (52) | 6.32 ± 0.38 (113) | 3371.00, 0.129 | 0.82, 0.286 |
| Information rate | 0.85 ± 0.11 (52) | 0.61 ± 0.05 (113) | 3387.00, 0.116 | 0.11, 0.134 |
| Interspike interval cv | 2.38 ± 0.12 (52) | 2.36 ± 0.09 (113) | 2996.00, 0.840 | 0.02, 0.904 |
| In-field mean rate | 3.78 ± 0.39 (43) | 2.54 ± 0.15 (100) | 2813.00, 0.004 | 0.94, 0.008 |
| Out-field mean rate | 0.34 ± 0.05 (52) | 0.50 ± 0.05 (113) | 2117.00, 0.004 | 0.15, 0.009 |
| Burst event ratio | 0.15 ± 0.02 (52) | 0.19 ± 0.01 (113) | 2413.50, 0.066 | 0.03, 0.165 |
| Specificity | 1.05 ± 0.06 (43) | 0.77 ± 0.03 (100) | 3051.00, 0.000 | 0.28, 0.000 |
| Speed score | 0.04 ± 0.01 (52) | 0.03 ± 0.01 (113) | 3128.00, 0.506 | 0.01, 0.835 |
| Fields area mean | 1152.99 ± 24.42 (52) | 1061.76 ± 13.05 (113) | 3812.00, 0.002 | 60.90, 0.011 |

**Figure S1:**
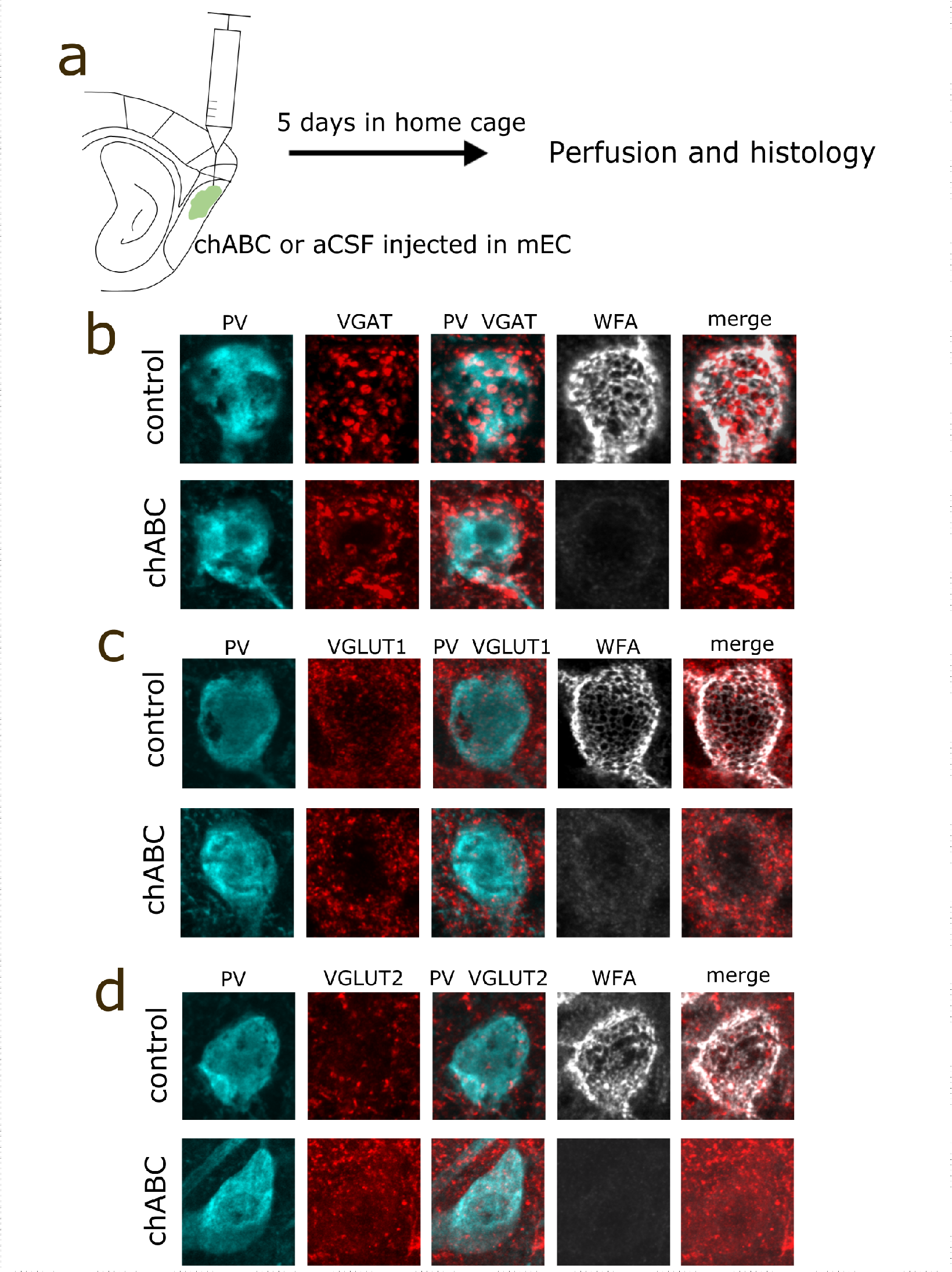
Synaptic bouton analysis after PNN removal. **a**) Experimental overview. chABC and aCSF was injected into MEC of one hemisphere each. After five days in the home cage, animals were perfused and sections from both hemispheres stained and imaged in parallel for parvalbumin, WFA and presynaptic markers. **b-d**) Example images from aCSF and chABC treated MEC tissue.

**Figure S2:**
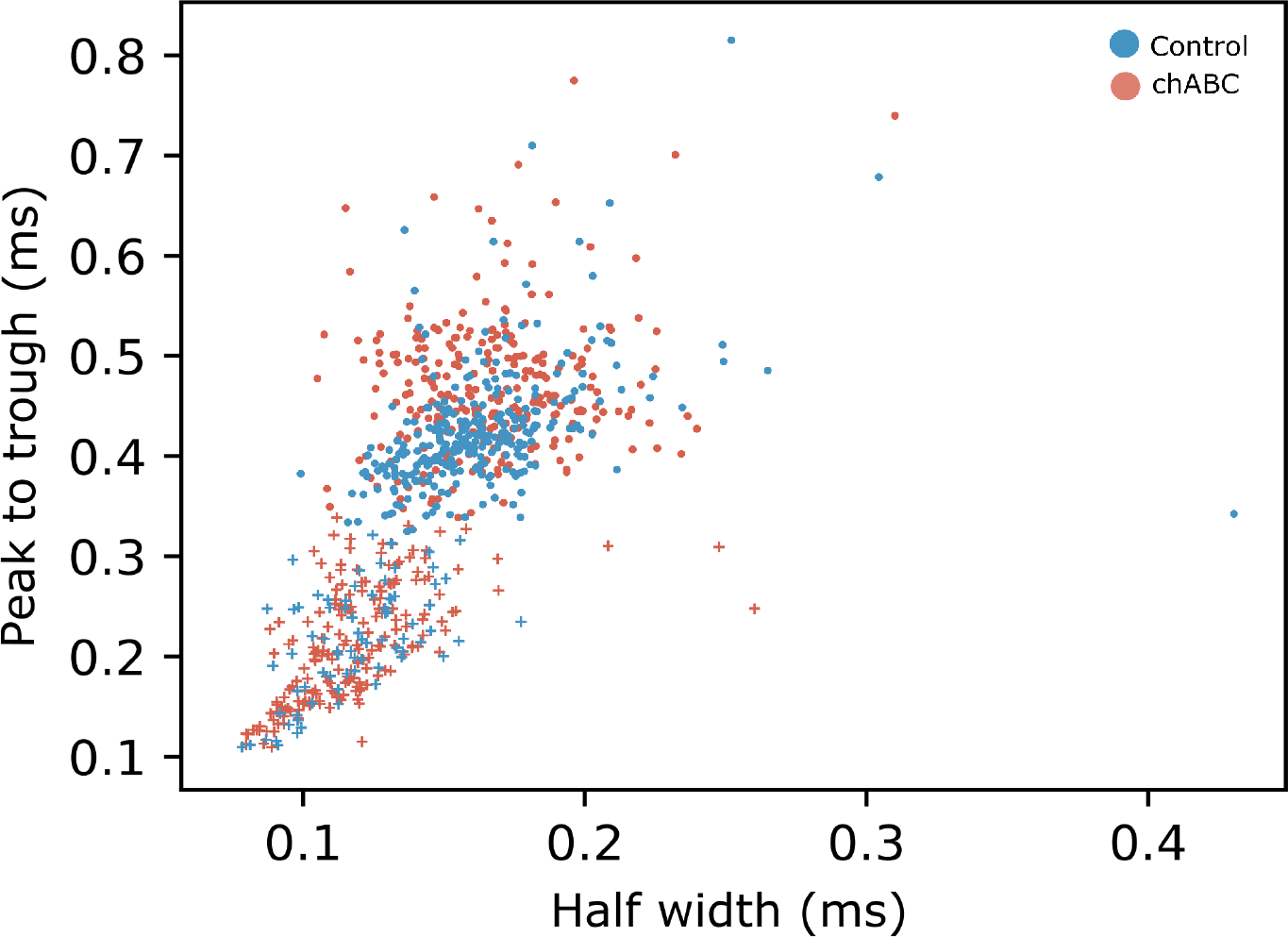
Separation of broad and narrow spiking units. Units are separated into broad or narrow spiking based by plotting peak to trough time against the width or half amplitude using kmeans. Circles represent units categorized as broad spiking and cross represent units categorized as narrow spiking.

**Figure S3:**
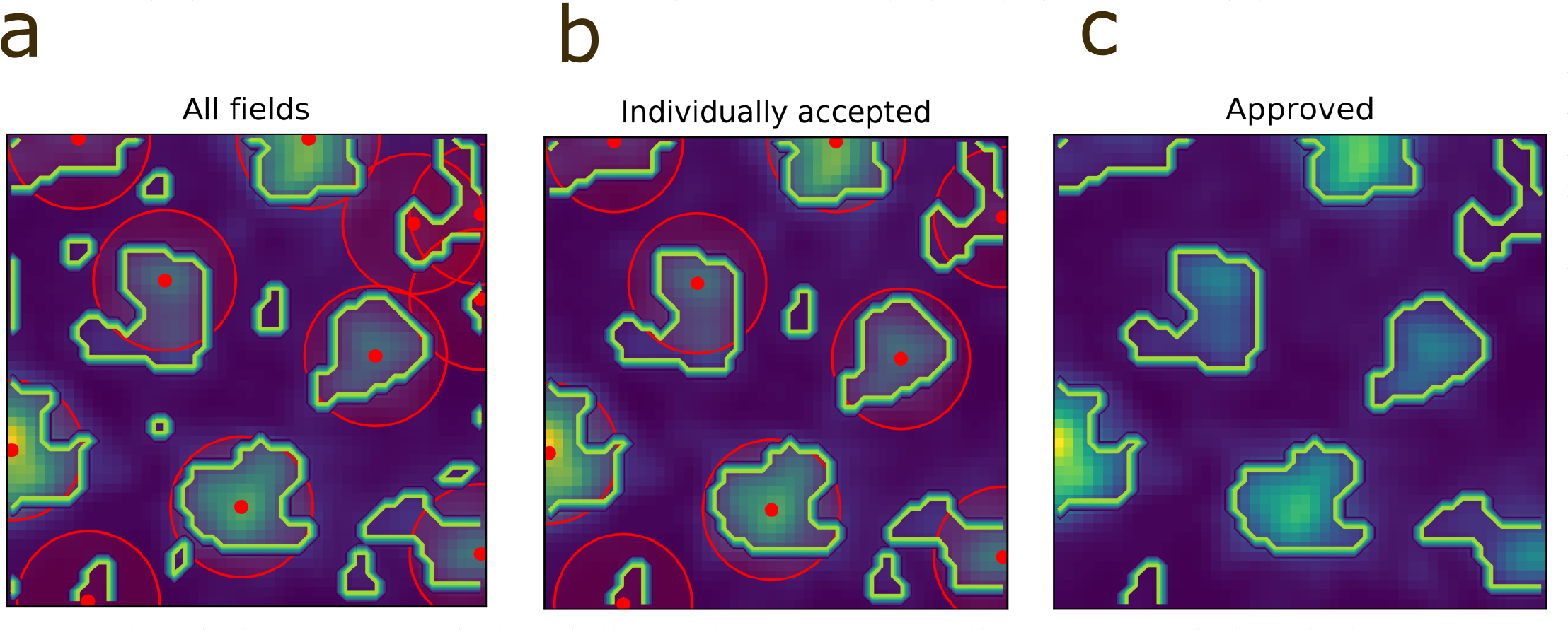
Grid field analysis. **a**) To estimate the firing within and outside fields, we first identified the individual fields in the rate map using a dual methods approach. First we defined the extent of each field, by calculating the Laplacian ∇^2^ of the smoothed rate map to obtain its curvature (outlined by blue and yellow). Regions with a positive Laplacian, which are the valleys of the rate map, were excluded. To limit random out-of-field rate getting considered individual fields using the laplacian, we included the protocol of [Ismakov et al., 2017], where we calculated a global field radius using 0, 7 times half the distance from the center peak to the closest peak in the autocorrelogram (field center is marked by a red dot and the global field radius is marked by a red circle). **b**) Next, we excluded the lowest of any two peaks within a distance shorter than the global field radius along with any regions with an area less than 9 bins. **c**) Fields were defined to be any labeled region found after taking the Laplacian that corresponded with a non-exluded peak from the protocol of [Ismakov et al., 2017].

**Figure S4:**
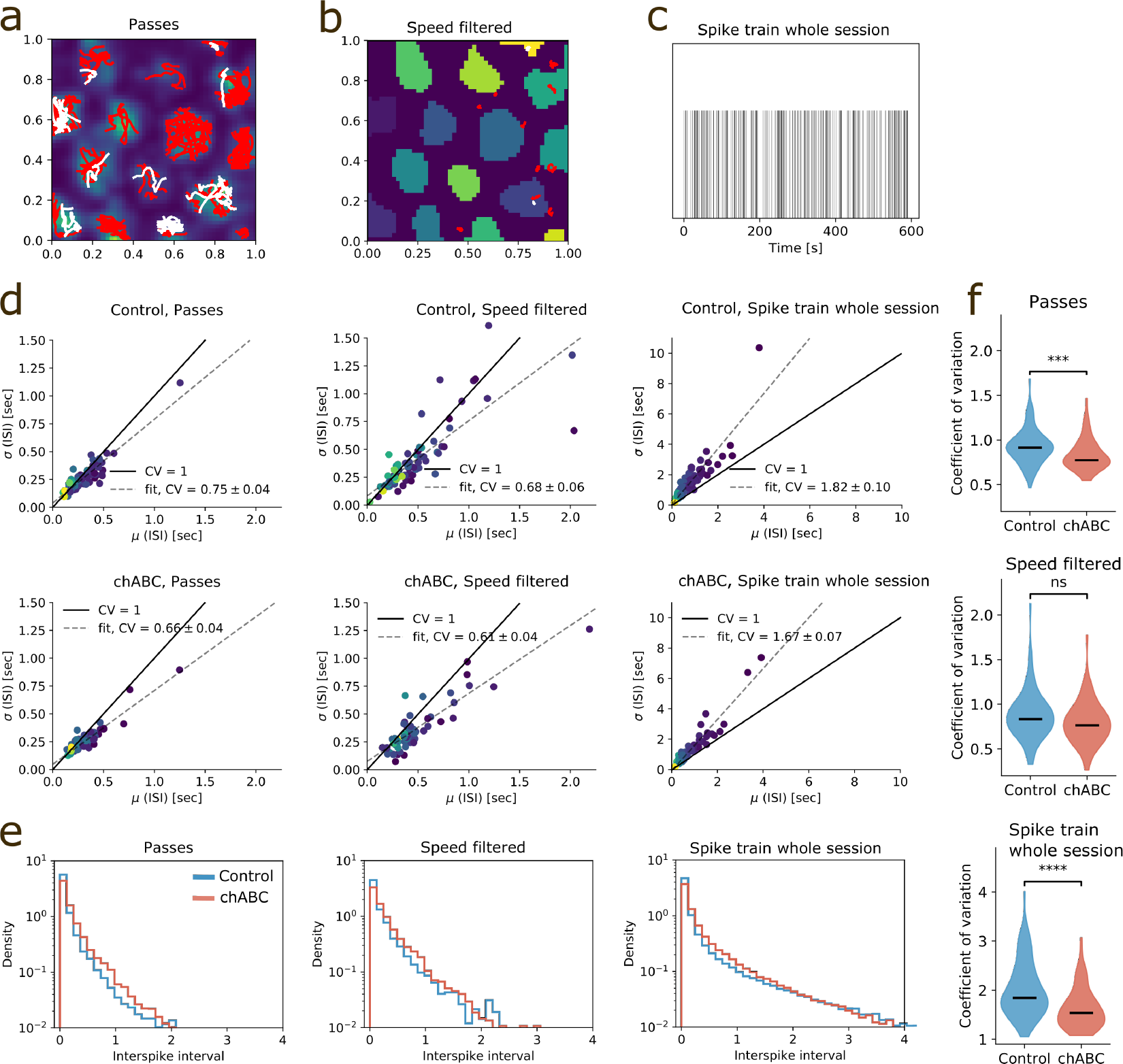
Three different methods for calculating CV of interspike interval for grid cells in a familiar environment. **a**) Passes through fields identified as described in Fig. S3. ISIs from all passes with duration longer than 1s and more than two spikes are used to calculate an ISI mean and standard deviation for the session. White (red) lines mark accepted (discarded) passes. **b**) Parts of the recording where the speed of the rat is lower than 8 cm/s, as described in (*60*). ISIs from all parts with duration longer than 1s and more than two spikes are used to calculate an ISI mean and standard deviation for the session. White (red) lines mark accepted (discarded) parts. White (red) lines mark accepted (discarded) parts. **c**) The 10 first minutes of a spike train. The Units method involves using the spike train from the whole session to calculate its ISI mean and standard deviation. **d**) Standard deviation of the interspike interval (ISI) *σ* versus mean ISI *μ*, for each of the three methods and both groups. Each data point is from a unique session, one session per unit. Groupwise CV is indicated by a least squares fit. Note the different axis limits due to larger values with the Units method. **e**) Distribution of the interspike interval for the three methods. Deviation from a straight line indicates deviation from a poisson process. **f**) Distribution of coefficient of variation for all three methods and both groups. Burak & Fiete: Control 0.88 ± 0.03 vs chABC 0.80 ± 0.03, *p* < 0.055. Passes: Control 0.93±0.06 vs chABC 1.64±0.05, *p* < 0.001. Units: Control 2.02±0.06 vs chABC 1.64±0.05, *p* < 0.001.

**Figure S5:**
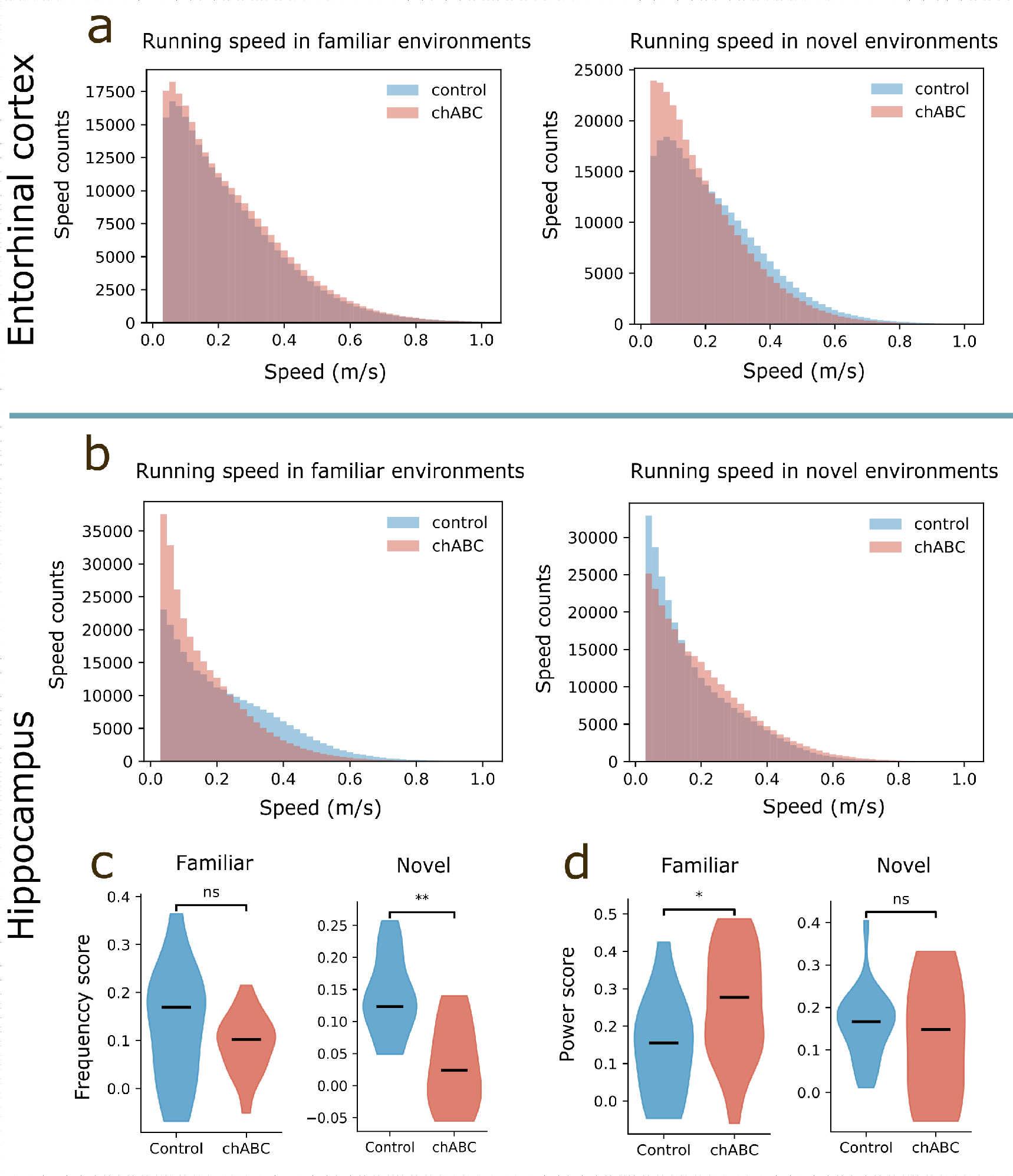
Running speed for all animals during exploration of a familiar and a novel arena. **a**) Running speed from all animals where we recorded local field potential (LFP) in MEC (upper panel) and **b**) hippocampus (lower panel). **c**) Frequency score (the correlation between instantaneous running speed and theta frequency) from recordings of LFP in hippocampus in familiar and novel environments. **d** Power score (the correlation between instantaneous running speed and theta power) from recordings of LFP in hippocampus in familiar and novel environments

**Figure S6:**
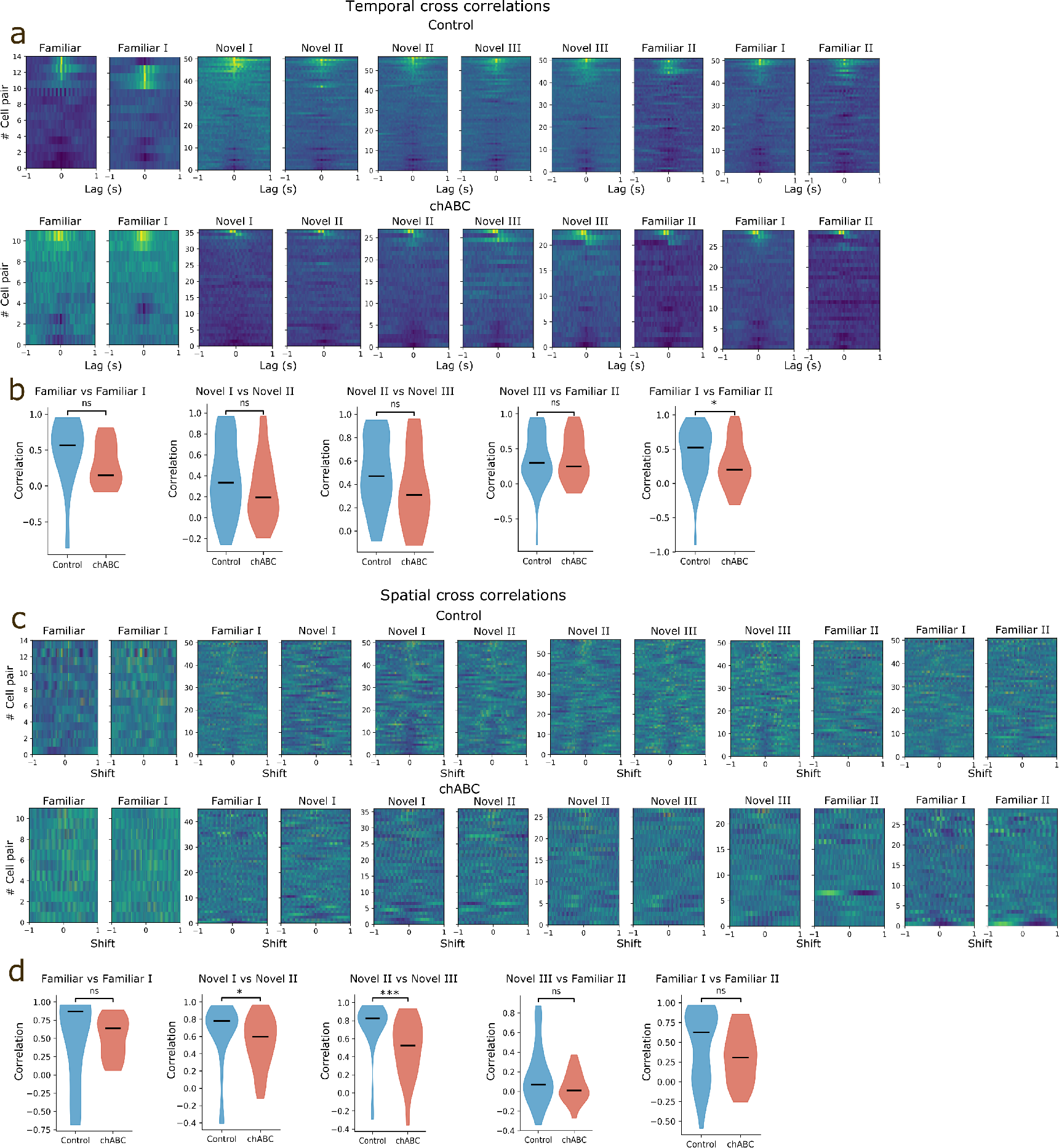
Pairwise correlations for all environments. **a**) Pairwise temporal correlation histograms of grid cells with brighter color showing zscored number of co-occuring spikes. Each line represents a pair, sorted by maximum value of central peak, with pair identity maintained across experimental states (left and right panel) **b**) Pairwise temporal correlations quantified for all environments. Data for Familiar I vs Novel I is shown in main figure. Familiar I includes all units recorded in the familiar environment on the first day of novel environment experiment. Familiar includes all units form any day that could also be identified in Familiar I. Familiar vs Familiar I; Control 0.49±0.13(*n* = 14); chABC 0.30±0.09(*n* = 11), *U* = 106.00*,p* = 0.119, Novel I vs Novel II; Control 0.37±0.05(*n* = 51); chABC 0.26±0.05(*n* = 36), *U* = 1080.00,p = 0.045, Novel II vs Novel III; Control 0.49±0.04(*n* = 57); chABC 0.36±0.06(*n* = 27), *U* = 960.00,p = 0.069, Novel III vs Familiar II; Control 0.36±0.05(*n* = 51); chABC 0.36±0.07(*n* = 23), *U* = 587.00*,p* = 1.000. Familiar I vs Familiar II; Control 0.43±0.05(*n* = 51); chABC 0.27±0.06(*n* = 29), *U* = 950.00, *p* = 0.036 **c** Figure organized as in **a**) showing spatial pairwise correlations.**d** Pairwise spatial correlations were significantly reduced in chABC animals in the novel environment, likely reflecting the slow stabilization of new grid fields in the chABC group. Familiar vs Familiar I; Control 0.48±0.17(*n* = 14); chABC 0.51±0.08(*n* = 11), *U* = 97.00*,p* = 0.286, Familiar I vs Novel I; Control 0.28±0.05(*n* = 51); chABC 0.03±0.03(*n* = 45), *U* = 1495.00,p = 0.011, Novel I vs Novel II; Control 0.64±0.05(*n* = 51); chABC 0.53±0.05(*n* = 36), *U* = 1216.00,p = 0.010, Novel II vs Novel III Control 0.74±0.04(*n* = 57); chABC 0.50±0.06(*n* = 27), *U* = 1195.00, *p* < 0.001, Novel III vs Familiar II; Control 0.16±0.04(*n* = 51); chABC 0.04±0.03(*n* = 23), *U* = 708.00,p = 0.158 Familiar I vs Familiar II; Control 0.46±0.06(*n* = 51); chABC 0.32±0.06(*n* = 29), *U* = 924.00, *p* = 0.066, Mann-Whitney U test.

**Figure S7:**
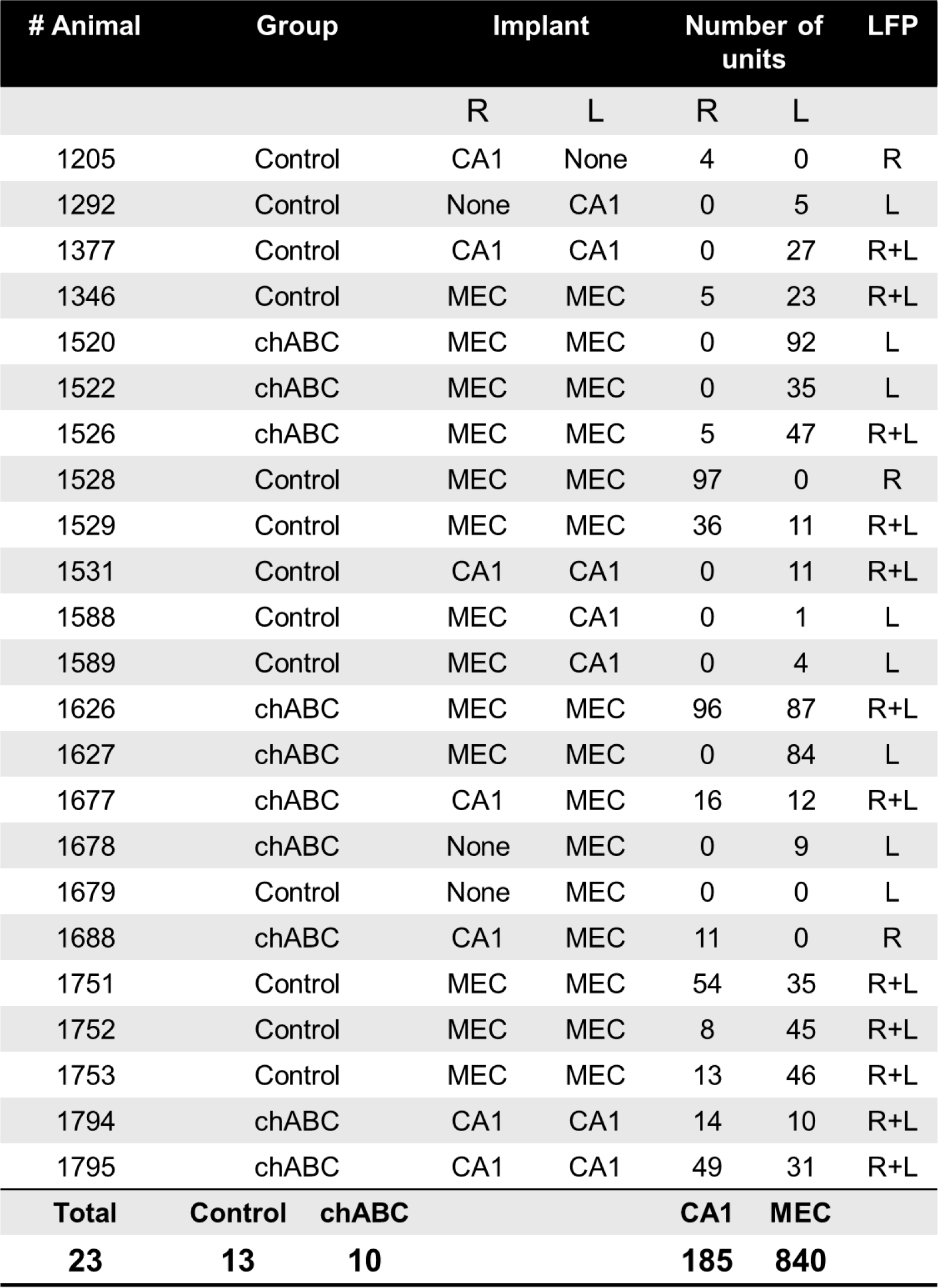
Overview of animals used for electrophysiological recordings.

**Figure S8:**
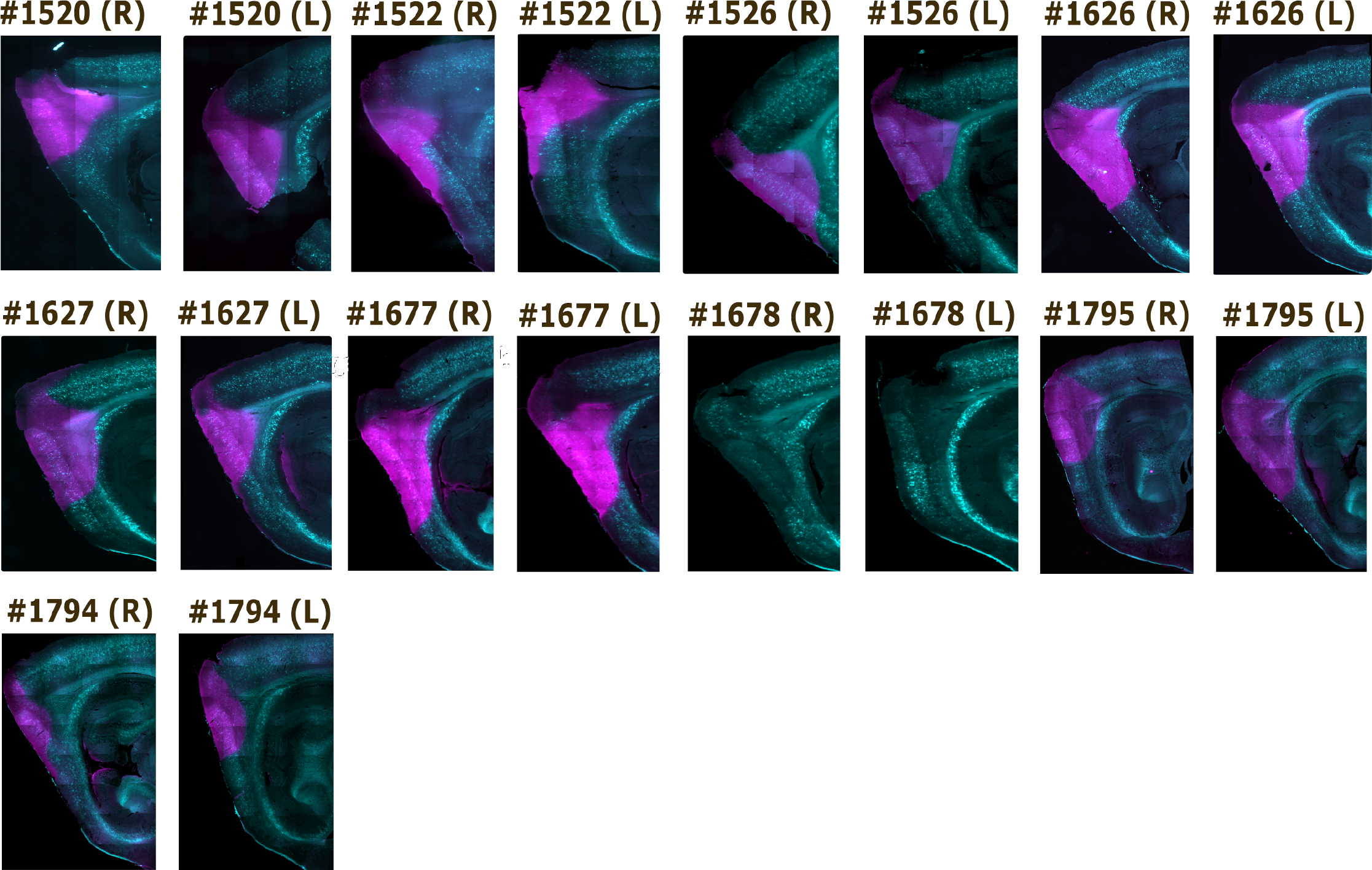
Confirmed PNN removal in MEC of experimental animals. Sections were stained with WFA (cyan) and 3B3 (magenta) that labels CS6 “stubs” left after chABC degradation of chondroitin sulfate proteoglycans. Note that animal #1678 is missing the 3B3 staining but the chABC injection is still clearly visible due to low intensity WFA staining in the injected area.

**Figure S9:**
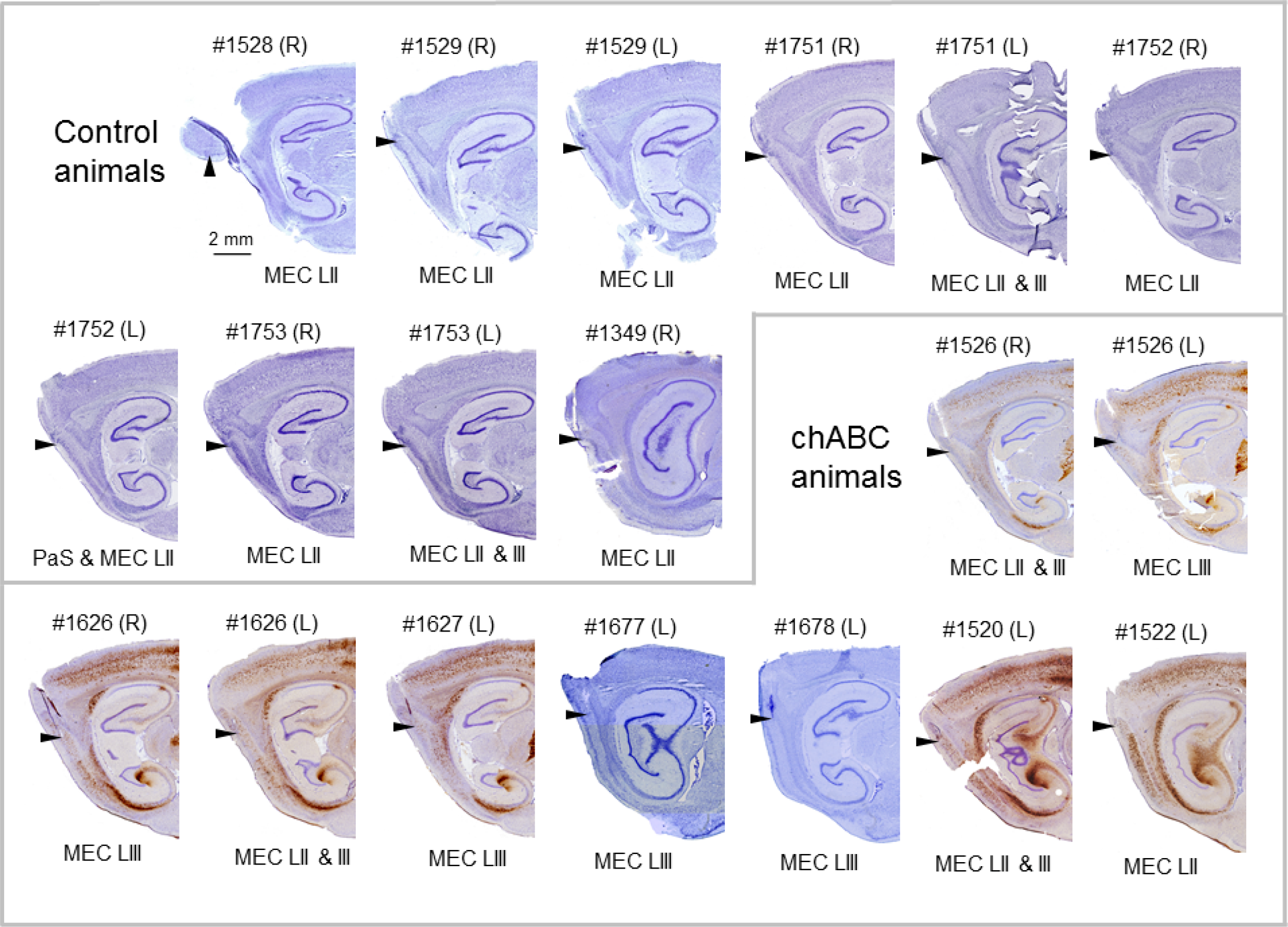
Tetrode tracks in MEC from experimental animals. Sections were stained for Nissl bodies to visualize the displacement of cell somas, and WFA labeled with an avidin-streptavidin - 3,3 diaminobenzidine reaction to visualize the area with PNNs removed. Arrows indicate the end of tetrode tracks, numbers indicate animal number, and (R) and (L) indicate the hemisphere with the brain seen from above in the anterior-posterior direction.

**Figure S10:**
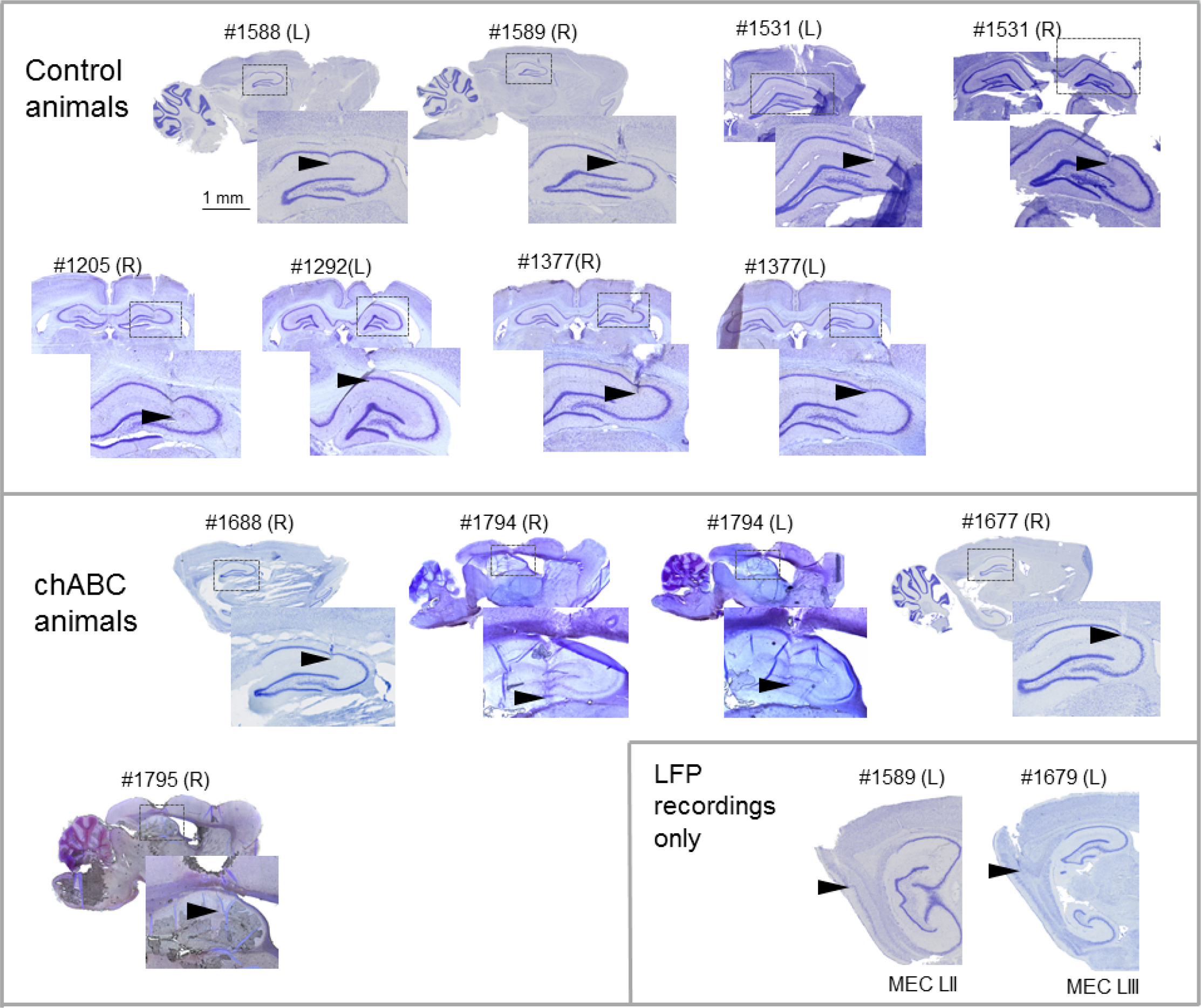
Tetrode tracks in hippocampus from experimental animals. Sections were stained for Nissl bodies to visualize the displacement of cell somas. Arrows indicate the end of tetrode tracks, numbers indicate animal number, and (R) and (L) indicate the hemisphere with the brain seen from above in the anterior-posterior direction. Lower right inset shows tetrode tracks in MEC from the left hemisphere of two control animals where we included animals in analysis of LFP recordings but did not record any units.

## Footnotes

1 neuralensemble.org/elephant

2 http://neuralensemble.org/elephant/

3 https://www.astropy.org/

4 github.com/regeirk/pycwt

